# Community Structure and Function During Periods of High Performance and System Upset in a Full-Scale Mixed Microalgal Wastewater Resource Recovery Facility

**DOI:** 10.1101/2024.01.23.576871

**Authors:** Md Mahbubul Alam, Mahdi Hodaei, Elaine Hartnett, Benjamin Gincley, Farhan Khan, Ga-Yeong Kim, Ameet J. Pinto, Ian M. Bradley

## Abstract

Microalgae have the potential to exceed current nutrient recovery limits from wastewater, enabling water resource recovery facilities (WRRFs) to achieve increasingly stringent effluent permits. The use of photobioreactors (PBRs) and the separation of hydraulic retention and solids residence time (HRT/SRT) further enables increased biomass in a reduced physical footprint while allowing operational parameters (e.g., SRT) to select for desired functional communities. However, as algal technology transitions to full-scale, there is a need to understand the effect of operational and environmental parameters on complex microbial dynamics among mixotrophic microalgae, bacterial groups, and pests (i.e., grazers and pathogens) and to implement robust process controls for stable long-term performance. Here, we examine the first full-scale, intensive WRRF utilizing mixed microalgal for tertiary treatment in the US (EcoRecover, Clearas Water Recovery Inc.) during a nine-month monitoring campaign. We investigated the temporal variations in microbial community structure (18S and 16S rRNA genes), which revealed that stable system performance of the EcoRecover system was marked by a low-diversity microalgal community (*D_INVSIMPSON_* = 2.01) dominated by *Scenedesmus* sp. (MRA = 55%-80%) that achieved strict nutrient removal (effluent TP < 0.04 mg·L-1) and steady biomass production (TSS_monthly avg._ = 400-700 mg·L^-1^). Operational variables including pH, alkalinity, and influent ammonium (NH_4_^+^), correlated positively (*p* < 0.05, method = Spearman) with algal community during stable performance. Further, the use of these parameters as operational controls along with N/P loading and SRT allowed for system recovery following upset events. Importantly, the presence or absence of bacterial nitrification did not directly impact algal system performance and overall nutrient recovery, but partial nitrification (potentially resulting from NO_2_^-^ accumulation) inhibited algal growth and should be considered during long-term operation. The microalgal communities were also adversely affected by zooplankton grazers (ciliates, rotifers) and fungal parasites (*Aphelidium*), particularly during periods of upset when algal cultures were experiencing culture turnover or stress conditions (e.g., nitrogen limitation, elevated temperature). Overall, the active management of system operation in order to maintain healthy algal cultures and high biomass productivity can result in significant periods (>4 months) of stable system performance that achieve robust nutrient recovery, even in winter months in northern latitudes (WI, USA).

**Graphical abstract:** 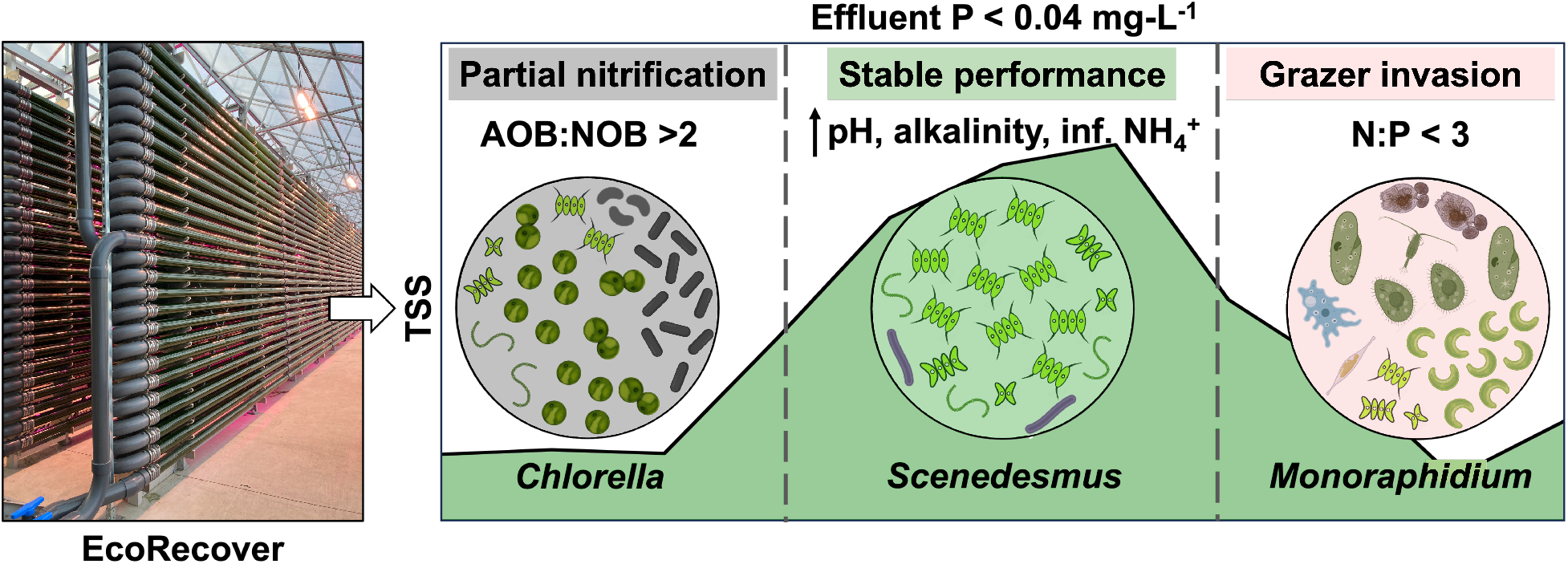

**Highlights:**

- Microbial dynamics were examined for first full-scale, intensive (small footprint) algal wastewater treatment process (EcoRecover) for advanced P removal.
- Mixed microbial communities during stable performance were dominated by *Scenedesmus* and Cyanobacteria and positively correlated with pH, alkalinity, and influent NH_4_^+^, among other parameters.
- Bacterial nitrification did not benefit or hinder nutrient recovery, but partial nitrification and NO_2_^-^ accumulation inhibited algal growth.
- Taxa specific pest dynamics are described, with major outbreaks occurring during high temperature in summer months.
- Control of operational parameters, and recovery of stable system performance and algal biomass was achieved following system upsets.

## 1. Introduction

Surface water discharge guidelines for wastewater nutrients (e.g., nitrogen (N) and phosphorus (P)) have become increasingly stringent (US EPA, 2023), especially in areas sensitive to eutrophication due to high agricultural and urban runoff (e.g., in the US Midwest and Chesapeake Bay) (Duan et al., 2012). Faced with these challenges, wastewater treatment utilities must implement sustainable and cost-effective technologies to achieve N and P removal below new limits (e.g., 0.5 mg-N·L^-1^ and 0.03 mg-P·L^-1^). Microalgae (i.e., unicellular eukaryotic phototrophs) have great potential to remove total N and P to levels below the current limit-of-technology (LOT) as well as the ability to utilize nutrients via assimilation which are otherwise unmitigated (e.g., organic N and P) in conventional treatment processes. In addition, algal biomass can be harvested and transformed into economically viable by-products such as biofuels (Moreira et al., 2023), biopolymers (Lutzu et al., 2021), and fertilizers (Ngoc-Dan Cao et al., 2023). Indeed, integration of algal biomass cultivation with wastewater treatment is critical for the economic and environmental sustainability of algal feedstock production (Kandasamy et al., 2023).

In contrast to monoculture algal cultivation, treatment of wastewater using microalgae leverages mixed bacterial and microalgal communities, which have been shown to achieve better nutrient removal, higher biomass productivities (Liu et al., 2017; Luo et al., 2020a), and increased community stability (Novoveská et al., 2016). In addition, mixed communities offer the considerable advantage of resilient cultures that may suffer community shift and some deterioration in performance, but not complete culture crash and loss of system function. However, understanding the complex impacts of operational, environmental, and biological factors on bacterial and algal dynamics to achieve uninterrupted nutrient recovery and biomass cultivation requires extensive knowledge about system drivers that are not well understood. Changing influent parameters and perturbations (Ding et al., 2021), imbalanced microbial communities (e.g., excessive nitrification) (González-Camejo et al., 2022), and pests (e.g., zooplankton grazers, fungal and bacterial parasites) (Molina-Grima et al., 2022) can all impact system performance and result in the loss of overall biomass productivity.

In full-scale wastewater treatment, mixed microalgae are most often cultivated in oxidation ponds or waste stabilization ponds (WSPs) as a byproduct, where they provide some oxygen for bacterial biochemical oxygen demand (BOD) removal and additional nutrient polishing (Craggs et al., 2003; Mahapatra et al., 2022). However, recent advancements include high-rate algal ponds (HRAPs) that provide better mixing and light exposure of cells, improving on the nutrient removal efficiencies and biomass productivity (Sutherland et al., 2017a; Sutherland and Ralph, 2020), and the separation of hydraulic retention and solids residence time (HRT and SRT) (Bradley et al., 2019; Luo et al., 2020b; Molitor et al., 2023) to significantly reduce the physical footprint of the system and achieve biomass concentrations 2-3.5× that of ponds for bioenergy production (Marbelia et al., 2014). The current knowledge of mixed algal-bacterial systems largely focuses on how operational and environmental parameters, including neutral pH (Sutherland et al., 2014) and high alkalinity supporting carbon uptake, short SRT (≤ 5 d) for nutrient assimilation (Bradley et al., 2019), and moderate temperature and light intensity (Gonzalez-Camejo et al., 2019; Meng et al., 2019) promote nutrient removal and biomass productivity.

Thus far, only a few studies have looked into the complex microbial dynamics within the mixed microalgal systems, including inter-domain interactions among mixotrophic microalgae, heterotrophic bacteria, autotrophic nitrifiers, and pests (Hoeger et al., 2022; Mattsson et al., 2021; Sutherland et al., 2017b), with most studies focusing on the interactions between algae and nitrifiers (e.g., (Krustok et al., 2016; Kwon et al., 2019)). Co-cultivating microalgae with nitrifying bacteria offers the advantage of microalgae supplying sufficient oxygen for full nitrification without mechanical aeration, while nitrification, in turn, reduces microalgal inhibition by free ammonia, especially when treating high strength wastewater (Casagli et al., 2021; Kwon et al., 2019). However, negative interactions between microalgae and nitrifiers can arise from their competition for resources (e.g., inorganic carbon) under certain operational conditions (e.g., high influent ammonium, temperatures, and light intensity) (González-Camejo et al., 2022). In addition, mixed microalgal systems are subject to the introduction of pests (e.g., zooplankton grazers) which can cause rapid declines in biomass productivity and nutrient removal (Montemezzani et al., 2016). Therefore, achieving and sustaining an efficient mixed microalgal consortia for full-scale wastewater treatment requires a comprehensive understanding of cross-domain microbial dynamics and their associated abiotic drivers.

To this end, this study investigated the temporal variations in microbial community structure and dynamics at the first full-scale, intensive (i.e., small footprint, high biomass) mixed-microalgal wastewater system in the US (EcoRecover; Clearas Water Recovery Inc.). The system was designed for tertiary wastewater treatment to remove phosphorus (P) below new permit limits at the Village of Roberts (VoR), WI of 0.04 mg-P·L^-1^. The facility consists of a full-scale (568 m^3^·day^-1^ or 0.15 MGD) plant with primary and secondary treatment, followed by mixed microalgal photobioreactors (PBRs) that leverage HRT/SRT separation for the rapid nutrient assimilation and removal of P via microalgal biomass. We hypothesized that microalgal and bacterial communities (including nitrifiers) would show significantly different dynamics during the steady-state operation and system upsets, and that these distinct periods of performance could be correlated with operational and environmental parameters. A nine month-long monitoring campaign was conducted and system performance, including water quality and biomass composition continuously monitored. Biomass was characterized weekly for microbial community analysis via amplicon sequencing of 16S and 18S rRNA genes for the identification of key organisms and their temporal dynamics. Correlation analysis was performed between microbial community data and physical and chemical data from the EcoRecover system to identify the fundamental drivers of microbial interactions and system performance. Ultimately, insights into operational and environmental controls and their effect on complex mixed-microalgal communities will aid in the adoption of robust, full-scale algal technology at WRRFs.

## 2. Materials and methods

### 2.1. EcoRecover system operation

For a detailed description of the EcoRecover system operation, refer to (Molitor et al., 2023). Briefly, the EcoRecover system contains three major operational units: a mix tank, PBRs, and membrane tank (Figure S1). The mix tank (design HRT: ∼ 3-4 h) receives secondary effluent from an upstream WRRF (0.15 MGD) at Village of Roberts (VoR), WI, USA and recycled algal sludge from the EcoRecover system. The majority (average 50-75%) of the ammonium and phosphate removal occurs in the mix tank under dark conditions as algae come into contact with nutrient-rich secondary effluent and deplete their carbon reserves (i.e., endogenous respiration). The mixed liquor is then sparged with CO_2_ and pumped through the PBRs, which consist of 10 parallel sets of PBRs (design HRT: ∼ 2-2.5 h). PBRs are located in a greenhouse utilizing natural sunlight supplemented with LED light (night-time; average: 45-77 µmol·m^-2^·s^-1^) to increase biomass yield. Algae accumulate storage compounds (e.g., lipid, carbohydrate) under phosphorus-limited conditions in the PBR. It is then pumped through the membrane tank (design HRT: < 30 min), where the separation of SRT (∼3-4 d) and HRT is achieved using two submerged ultrafiltration cassettes (0.03 µm). Finally, a portion of the permeate is stored for reuse, and membrane retentate, PBR recycle, and centrate are mixed in the return tank and recycled to the mix tank.

### 2.2. Long-term EcoRecover monitoring and biomass collection

On-line sensors were installed in the effluent streams of all three operational units (mix tank, PBR, and membrane tank) for long-term monitoring of operational parameters (e.g., pH, DO, temperature, total suspended solids (TSS), turbidity, flow, photosynthetically active radiation (PAR), and nutrients (PO_4_^3-^, NH_4_^+^ NO_3_^-^). Real-time data from the on-line sensors were acquired through a supervisory control and data acquisition (SCADA) system. In addition, grab sampling was performed one to two times a day from each unit to complement on-line SCADA data. Aqueous water parameters (PO_4_^3-^, total phosphorus, NH_4_^+^, NO_2_^-^, NO_3_^-^, total nitrogen, pH, alkalinity) and growth parameters (TSS, VSS) were measured using Standard Methods (Table S1). Data cleaning and smoothing were performed for each operational parameter for the periods when SCADA system was down or showed unreliable data due to malfunctions in on-line sensors (Molitor et al., 2023).

In parallel with long-term monitoring and system characterization, biomass was collected weekly for nine months to monitor the changes in microbial community structure over time and their correlation with operational changes. 1 mL of suspended biomass was collected in triplicate from the mixed tank and PBR effluent and stored in 5 mL polypropylene transport tubes (Fisherbrand^TM^, USA) prefilled with 3 mL DNA/RNA Shield^TM^ (Zymo Research corp., CA, USA). The samples were briefly vortexed and stored at -20 °C until they were shipped to University at Buffalo (UB) SUNY (Buffalo, NY) overnight on ice.

### 2.3. DNA extraction, PCR, and amplicon sequencing

DNA extraction was performed in duplicate using DNeasy PowerSoil Pro Kit (QIAGEN, USA) according to the manufacturer protocol. DNA extracts were immediately stored at -20 ℃ before further analyses. The concentrations of DNA were measured using Qubit^TM^ dsDNA High Sensitivity (HS) Assay kit (Invitrogen^TM^) via Qubit^TM^ 2.0 fluorometer (Invitrogen^TM^). After DNA extraction, the 18S rRNA gene was amplified by targeting the V8-V9 variable region [forward V8f: 5’-ATAACAGGTCTGTGATGCCCT-3’; reverse V9R: 5’-CCTTCYGCAGGTTCACCTAC-3’ developed by (Bradley et al., 2016)], and 16S rRNA gene was amplified by targeting the V4 region (forward 515F: 5′-GTGYCAGCMGCCGCGGTAA-3′, reverse [806R] = 5′-GGACTACNVGGGTWTCTAAT-3′) (Apprill et al., 2015; Parada et al., 2016). Full-length primers containing the adapters, index, and pad/linkers for Illumina MiSeq sequencing were designed according to the dual-index method (Kozich et al., 2013) and purchased from Integrated DNA Technologies (Coralville, IA, USA). The PCR amplification was performed in two technical replicates using KAPA Hifi Hotstart PCR kit (Kapa Biosystems, MA, USA) via a Bio-Rad CFX96 thermal cycler (Bio-Rad Laboratories) with reagent concentrations and thermocycler conditions for V8-V9 (18S rRNA) and V4 (16S rRNA) assays as described previously by (Bradley et al., 2016) and (Alam et al., 2022), respectively. A negative PCR control was included for each primer set to ensure absence of contamination during amplification. Following PCR amplification, gel electrophoresis was performed on all amplicon samples to excise the bands of expected size and quality (V8-V9 = 490 bp, V4 = 390 bp). The excised gel bands were extracted and purified using QIAquick gel purification kit (QIAGEN, USA). The concentrations of the purified DNA amplicons from each sample were measured in triplicates (as previously described) and pooled in equimolar proportions (∼10-20 ng) into two separate libraries for 18S rRNA and 16 rRNA gene sequencing. The final amplicon libraries were sequenced using the Illumina MiSeq with v3 chemistry (300-cycle paired-end reads) on two separate runs (for 18S and 16S) at the UB Genomics and Bioinformatics Core. All raw sequencing data can be found online on NCBI under the BioProject accession number PRJNA930162.

### 2.4. Sequence read processing and statistical analyses

Sequence processing was performed using mothur v.1.48.0 following the standard operating procedure (MiSeq SOP; (Kozich et al., 2013)). Contigs were formed using forward and reverse reads followed by quality trimming of ambiguous base calls, merging duplicate reads, alignment with SILVA SSU v138.1 reference database (Quast et al., 2013; Yilmaz et al., 2014), pre-clustering and the classification of amplicon sequence variants (ASVs), and chimera detection and removal using the VSEARCH algorithm (Rognes et al., 2016). Sequences were then classified using SILVA SSU v138.1 database. To identify the unclassified ASVs at genus level, representative sequences from each ASV were extracted using the *get.oturep* command in mothur and matched against the NCBI’s nucleotide BLAST database for high sequence similarity (Madden, 2003).

All the statistical analyses associated with the alpha and beta diversity calculations were performed using built-in mothur commands (Schloss et al., 2009) unless described otherwise. Alpha diversity was estimated by measuring observed ASVs (Sobs) and calculating inverse Simpson index (invSimpson) and nonparametric Shannon index (npShannon) using the *summary.single* command, and beta diversity was assessed from Bray-Curtis dissimilarity matrix using the *dist.shared* command. The dissimilarity between samples was visualized via principal component analysis (PCoA) and non-metric multidimensional scaling (NMDS) using the Bray-Curtis dissimilarity matrix. The statistical differences among the samples from stable and upset periods were determined by the analysis of molecular variance (AMOVA), a nonparametric analog of ANOVA, which determines if the dissimilarity between samples within each community clusters are significantly different (Excoffier et al., 1992; Huang et al., 2021). ASV correlations with the PCoA and NMDS axes were analyzed using *corr.axes* command (*p* < 0.05, method = Spearman) to reveal which ASVs were responsible for community shifts during the stable and upset periods. Correlations (*p* < 0.05, method = Spearman) of the metadata containing abiotic factors (i.e., water quality and operational parameters) from each sample with the PCoA and NMDS axes were also performed. The correlation coefficients obtained for ASVs and metadata for each axis were then mapped on top of the PCoA and NMDS plots. In order to obtain the relationship between the abiotic factors and ASV abundance, a co-occurrence network analysis was performed using CoNet v1.1.1 and visualized via Cytoscape 3.9.1 with a significance threshold of 0.05.

## 3. Results and Discussion

Long-term EcoRecover operation was marked by “stable” and “upset” periods. Stable system performance was defined as consistent biomass production (TSS_monthly avg._ = 400-700 mg·L^-1^) and effluent P below permit levels of 0.04 mg-P·L^-1^ and was characterized by neutral pH, adequate alkalinity (≥ 200 mg·L^-1^), and a consistent daily DO cycling, while upset periods were typically defined by the loss of biomass productivity (TSS < 250 mg·L^-1^) and system performance (i.e., effluent TP (monthly avg.) > 0.04 mg·L^-1^) (Figure 1, Figure S2). Process operation during long-term monitoring experienced three distinct periods: (1) upset 1: a process upset due to partial nitrification and subsequent recovery (2021-11-04 to 2022-01-03); (2) stable performance: long-term stable operation (2022-01-04 to 2022-05-23); and (3) upset 2: a process upset due to sudden upstream influent changes (2022-05-24 to 2022-08-02).

**Figure 1.**
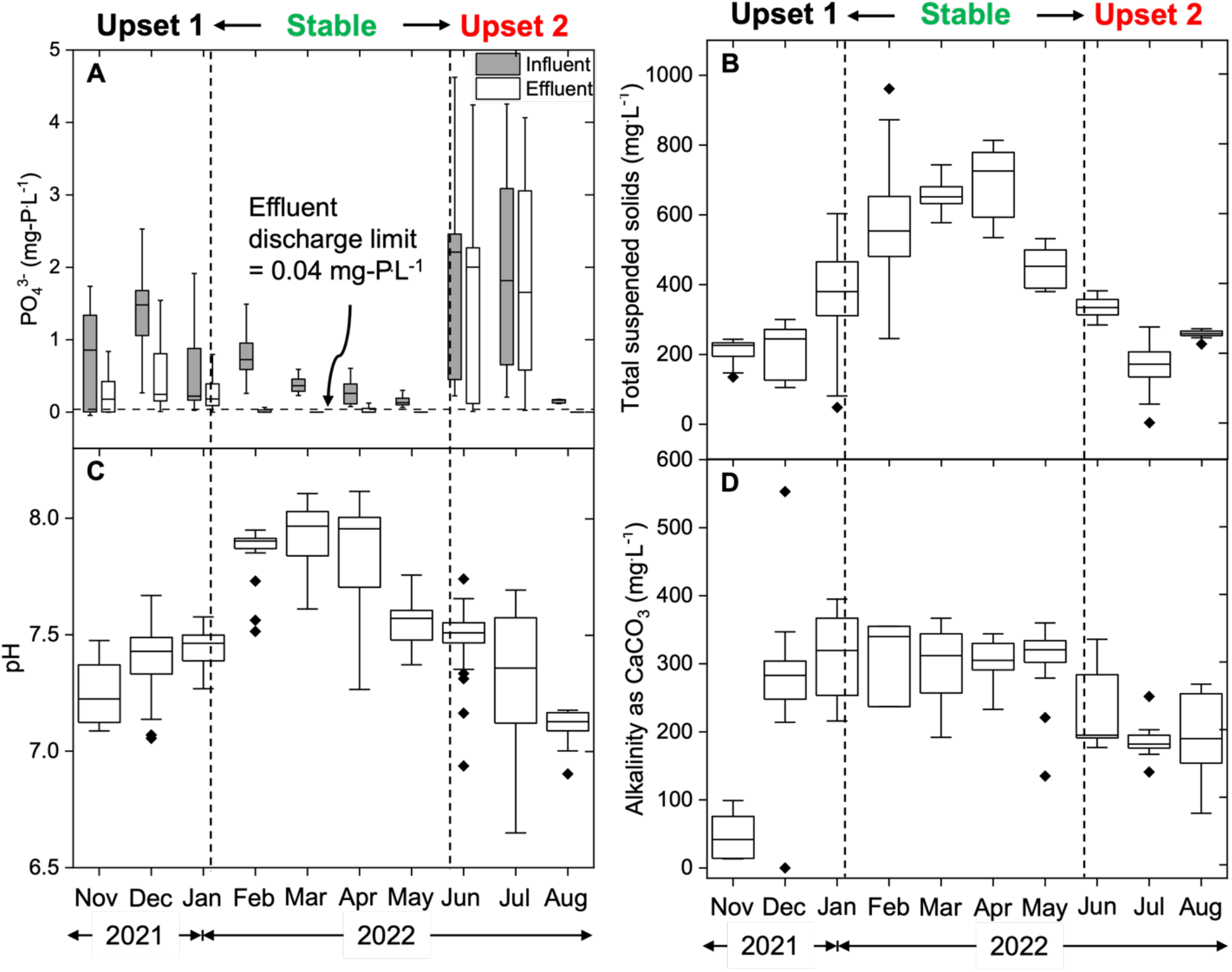
Average monthly operational parameters of the EcoRecover process for the period between November 4, 2021 to August 2, 2022: (A) influent and effluent orthophosphate (PO_4_^3-^) concentrations, (B) total suspended solids (TSS) of the mixed system, influent (C) pH and (D) alkalinity. Stable performance consistently met the effluent discharge permit (< 0.04 mg·L^-1^) with TSS ranging from 400-700 mg·L^-1^ under increased pH (7.5-8) and stable alkalinity 280-315 mg·L^-1^.

### 3.1. Long-term community structure and dynamics

Stable system operation (effluent TP_monthly avg._ < 0.04 mg·L^-1^, TSS_monthly avg._ = 400-700 mg·L^-1^; Figure 1) was characterized by high mean relative abundance (MRA; 55-80%) of *Scenedesmus* sp. with a significantly different (*p* < 0.01, AMOVA) community structure and lower eukaryotic diversity than upset periods (Figure 2). The eukaryotic communities were dominated by green microalgae (i.e., Chlorophytes; 52-97%), irrespective of the system performance (Figure 2A), with the most dominant species belonging to the genus *Scenedesmus, Chlorella*, *Monoraphidium*, and *Desmodesmus*. During the first upset period, *Chlorella* (52%) and *Desmodesmus* (39%) were the dominant genera, with *Chlorella* increasing to 77% and *Desmodesmus* decreasing to 19% as system operation continued to experience low nutrient uptake and biomass loss. Following significant operational changes to increase biomass productivity (further described in **section 3.3.1**) along with supplementation of algae paste (dominated by *Scenedesmus* sp.) from another EcoRecover system (Figure S3), a rapid turnover in the dominant species occurred, with *Scenedesmus* taking over as system performance started to improve. During the subsequent stable performance period lasting over ∼4.5 months, *Scenedesmus* dominated the eukaryotic community with a high mean relative abundance (MRA) of 55-80% while *Chlorella* became a minor constituent (0.5-13%) until it was no longer detected in the system. *Desmodesmus* continued to be the second-most abundant genus (10-30%) throughout the stable period. As spring advanced (March-early May, 2022), *Scenedesmus* reduced from 80% to 57%, while *Monoraphidium* increased almost 20 times from 0.4% to 8.5%. The onset of summer (May/June 2022) triggered upset period 2 which coincided with a change in upstream influent source (see **section 3.3.2**), switching the system from P-limitation to N-limitation, coupled with an increase in influent water temperature. During this time, another rapid species turnover was observed with a significant increase in the relative abundance of *Monoraphidium* (MRA > 70%) within a week, accompanied by a drastic decrease in the relative abundance of *Scenedesmus* (from 57% to < 1%). With the process recovery (late summer 2022), the relative abundance of *Scenedesmus* increased from 0.2% to 50%. The alpha diversity (species richness and abundance) indicated that the stable microalgal communities were less diverse than those during upset 1 and 2 periods [*D_INVSIMPSON_ (avg)* = 2.01 compared to 3.04 and 4.50, respectively (Figure 2C), *p* < 0.05, ANOVA and Tukey’s HSD], while the observed number of amplicon sequence variants (ASVs) was not significantly different during the stable period compared to upset periods [*D_sobs_ (avg)* = 134, versus 153 and 174 during upset 1 and 2 periods (Figure S4A), *p* > 0.05, ANOVA and Tukey’s HSD].

**Figure 2.**
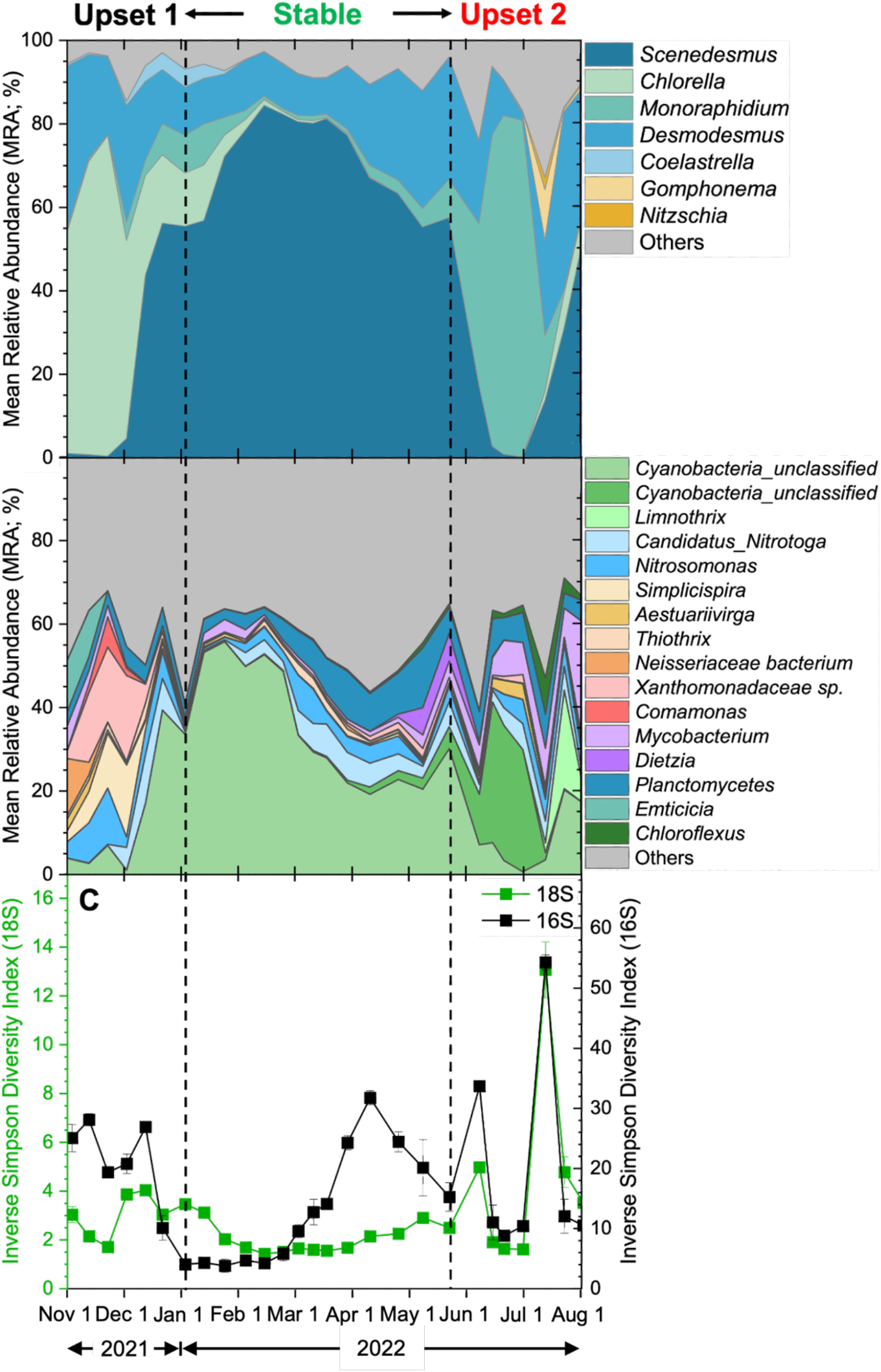
Mean relative abundance (MRA) of the top (≥ 1%) (A) microalgal and (B) bacterial genera derived from 18S rRNA and 16S rRNA amplicon gene sequencing, respectively. *Scenedesmus sp.* dominated the microalgal community during stable system performance which was characterized by high biomass productivity and nutrient removal (Figure 1), while upset periods had significantly (*p* < 0.01, AMOVA) different community structure and loss of biomass productivity. (C) Inverse Simpson diversity index of the eukaryotic (green) and bacterial (black) community. A comparatively less diverse eukaryotic community was complemented by a more diverse bacterial community.

Similar to the eukaryotic microalgal community, the bacterial community structure exhibited significant differences (*p* < 0.01, AMOVA) between stable performance and upset periods, with Cyanobacteria (30-56%) dominating during stable operation (Figure 2B). The bacterial community showed 5-7x higher diversity and species richness than the eukaryotic community (Figure 2C, Figure S4). The onset of the first upset period resulted in the proliferation of diverse facultative anaerobes (e.g., *Simplicispira*, *Emticicia*, *Comamonas* and multiple ASVs from Xanthomonadaceae family) and was subsequently characterized by increased abundance of nitrifying bacteria including *Nitrosomonas*-like and *Candidatus* Nitrotoga-like ASVs likely due to operational changes including increased SRT for stable nitrification (detailed in **section 3.3.1**). The initial stable bacterial community was primarily composed of Cyanobacteria ASV (50-56%, closely identified as *Arthrospira* on NCBI BLAST), but as the stable period progressed, additional bacterial taxa, such as *Planctomyces*, *Dietzia*, and *Mycobacterium* emerged in the system (Figure 2B). During the second upset period, a cyanobacterial ASV (closely matched with *Loriellopsis* on NCBI BLAST) exhibited increased relative abundance (29-34%), while the MRA of *Arthrospira*-like ASV decreased from 30% to less than 1%. As the system started recovering from the second upset period, the *Arthrospira*-like ASV (recovered from 3.5% to 20%) and another Cyanobacteria genus *Limnothrix* (23%) co-dominated the community. Bacterial diversity during the first two months (January to early March 2022) of the stable period [*D_INVSIMPSON_ (avg)* = 4.6] was significantly lower compared to upset periods 1 and 2 [*D_INVSIMPSON_ (avg)* = 4.6, 19.2, 20.1, respectively, *p* < 0.05, ANOVA and Tukey’s HSD], but became more diverse in latter portion of the stable performance period [*D_INVSIMPSON_ (avg)* = 19.1]. The observed number of bacterial ASVs that contributed to 80% of the total MRA in each sample was significantly less (*p* < 0.05, ANOVA and Tukey’s HSD) during upset 1 compared to stable and upset 2 [*D_sobs_ (avg)* = 58, 103, 116 during upset 1, stable, and upset 2, respectively) (Figure S4B)].

The key microalgal genera observed in this study, i.e., *Scenedesmus*, *Chlorella*, *Monoraphidium*, and *Desmodesmus* have been frequently reported to have a higher biomass yield and better biochemical compositions in mixotrophic cultivation and in the presence of mutualistic wastewater bacteria (Divya Kuravi and Venkata Mohan, 2022; Song et al., 2021). In addition to the exchange of O_2_ and CO_2_ for a balanced aerobic respiration and photosynthesis, microalgae growth promoting bacteria (MGPB) (e.g., *Planctomyces* and *Mycobacterium*) observed in this study have been reported to produce cofactors (e.g., vitamins) and degradation products, signaling molecules (e.g., phytohormones), and organic pigments (e.g., carotenoid) that benefit algal growth (Ludington et al., 2017; Luo et al., 2014; Palacios et al., 2022). While it is commonly regarded that a consortium of diverse microalgal communities can demonstrate a broader functional spectrum and thus help to maintain stable and robust nutrient removal and biomass production (Cho et al., 2017), stable system performance here was marked by dominance of *Scenedesmus*, which may represent an efficient but potentially less resilient (in the absence of significant diversity) community structure (Bradley et al., 2019). Both *Scenedesmus* (55-80%) and *Desmodesmus* (10-30%) that were dominant during the stable operation period have been frequently reported in HRAP and PBR systems (Cho et al., 2017; Ferro et al., 2020) and are known for their high photosynthesis and nutrient removal capacity, fast growth, and high lipid and exopolysaccarides (EPS) accumulation efficiency (Silambarasan et al., 2023). Both *Scenedesmus* and *Desmodesmus* also have high phenotypic plasticity marked by the ability to form protective four- or eight-celled colonies that could give them a competitive advantage against environmental stressors, including grazing pressures from zooplanktons (Lürling, 2003) which were a small (≤ 6.5% MRA) but constant presence in the EcoRecover system (further described in **section 3.3.2)**. Particularly, in January 2022, an increase in colonial *Scenedesmus* population (four and eight-celled colonies) was observed via microscopy which was preceded by rise sharp rise in the relative abundance of ASVs within the phylum *Cercozoa* (primarily sequences belonging to the genus *Rhogostoma*, Figure S5) followed by a rise in the abundance of zooplankton grazer rotifers (2.3-3.5%) in the system (**section 3.3.2**). Morphological alterations in *Scenedesmus*, indicative of its self-defense mechanism, were noted even before the rotifer abundance surpassed the detection limit. Subsequently, the *Scenedesmus* population returned to a unicellular state as rotifers became nearly absent (< 0.1%) from the system.

### 3.2. Drivers of stable system performance

Eukaryotic community structure during the stable period were positively correlated with several key variables including system pH (*p* < 0.001, *r* = 0.57), alkalinity (*p* < 0.001, *r* = 0.50), and influent ammonium (NH_4_^+^; *p* < 0.01, *r* = 0.55) using Spearman correlations (Figure 3A). Non-metric multi-dimensional analyses (NMDS) conducted using Bray-Curtis dissimilarity of relative abundances showed that the eukaryotic communities during the stable period formed a distinct cluster that differed significantly (*p* < 0.01, AMOVA) from those observed during upset periods (Figure 3A). A biplot of abiotic variables generated using Spearman correlation coefficients and mapped over the NMDS axes revealed that the distances between the samples of the stable and upset period communities could be explained by a subset of abiotic factors, including a neutral to slightly basic pH ranging from 7.5 to 8, high alkalinity in the mixed system (280-315 mg·L^-1^ as CaCO_3_), elevated influent NH_4_^+^ levels (30-45 mg N·L^-1^), effluent NO_3_^-^ levels (10-30 mg N·L^-1^), and moderate to high light intensity (120-150 µmol·m^-2^·s^-1^; Figure 1, Figure S2). DO levels in the mix tank and PBR ranged from 1-10 mg·L^-1^. Influent and effluent OP concentrations varied from 0.15 to 0.8 mg P·L^-1^ and 0.01 to 0.03 mg P·L^-1^, respectively (monthly average < 0.04 mg·L^-1^). The influent N:P ratio throughout the stable period was kept above 25:1, while the system’s temperature was within the range of 12-18 °C.

**Figure 3.**
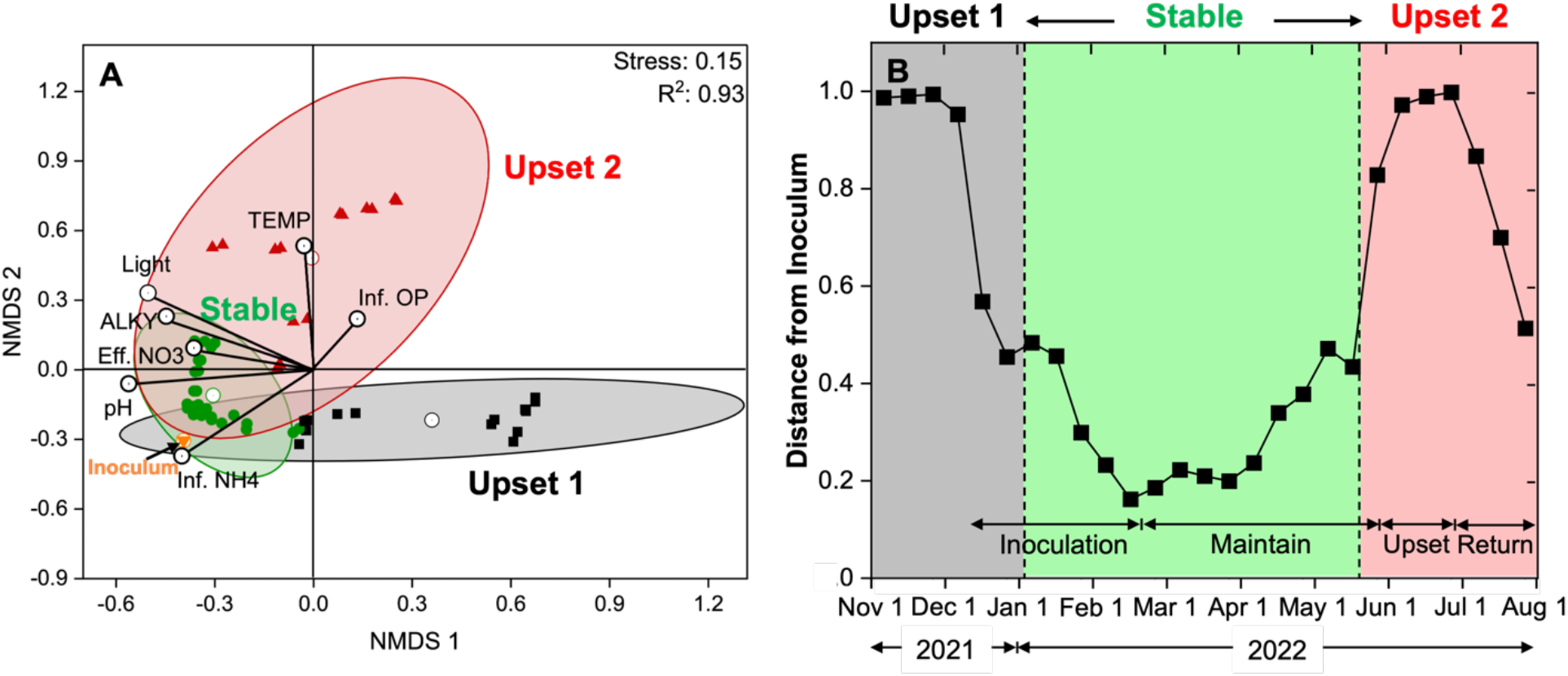
(A) Non-metric multi-dimensional scaling (NMDS) plot using Bray-Curtis dissimilarity showing distances among eukaryotic communities during periods of stable performance and upset events, with lines indicating strength and direction of correlation (method = Spearman) for each variable. Ellipses containing 95% cluster-assigned sampling points show that the community cluster during stable period was significantly different (*p* < 0.01, AMOVA) from upset periods and was associated with pH, alkalinity, and influent NH_4_^+^, as well as short distance from the inoculum culture. (B) Bray-Curtis distances of each sampling point from the inoculum community. The two upset periods showed significantly higher (*p* < 0.05, ANOVA and Tukey’s HSD) distance from the inoculum than the stable period and community recovery was observed at the end of the second upset period.

Maintaining beneficial operational parameters under changing environmental conditions is critical to ensure stable system performance in mixed community suspended systems (i.e., HRAPs, open ponds) (Cho et al., 2015; Ferro et al., 2020), and increased NH_4_^+^ has been previously associated with microalgal concentration in HRAPs (Cho et al., 2015) as it can more readily assimilate the reduced form of N. Although a high influent NH_4_^+^ concentration resulted in high bacterial nitrification (i.e., high effluent NO_3_^-^) in the EcoRecover system, a positive correlation was observed between high effluent NO_3_^-^ levels and stable eukaryotic communities (Figure 3A), indicating that microalgal growth remained unaffected when favorable conditions for microalgae (including pH, alkalinity, and light requirements) were met, despite competition from ammonia oxidizing bacteria (AOBs). In our study, in the middle of the first upset period (early December 2021), the system was supplied with sodium bicarbonate (NaHCO_3_) and ammonium (NH_4_^+^) (to maintain ≥ 200 mg·L^-1^ as CaCO_3_ and > 15 mg·L^-1^ NH_4_^+^-N, respectively) to increase alkalinity and pH levels and prevent ammonium limitation, respectively. This, along with supplementation of healthy algal biomass from early December to late February (comprising 6-20% of the total biomass and dominated by *Scenedesmus* sp.) shifted the community from a *Chlorella* to *Scenedesmus* dominated community. This approach allowed the maintenance of a culture similar to the inoculum (i.e., minimal community dissimilarity) for an extended period of over ∼4.5 months after the inoculation was stopped (Figure 3B). Additionally, a stable *Scenedesmus* community and return to P effluent discharge below permit levels were achieved following the second upset period by operational changes (e.g. N:P ratio) in the absence of further culture inoculation.

### 3.3. Drivers of system upset and unstable performance

Both upsets were characterized by substantial fluctuation in DO with inconsistent diel cycling, loss of productivity, and failure to meet monthly average effluent TP discharge limits of 0.04 mg·L^-1^. Each period was caused by a distinct event (Figure S2). The first upset period was caused by a loss of complete nitrification (< 20%) and nitrite (NO_2_^-^) accumulation that negatively impacted culture growth and system stability and was inversely correlated with pH and alkalinity. The second upset period occurred when a sudden switch in upstream loading transitioned the system from P to N limitation and was positively correlated (Figure 3A) with influent OP levels (1.9-2 mg·L^-1^), high system temperatures (22-25 °C), and subsequent increases in pest abundance.

#### 3.3.1. Upset 1 – Effect of bacterial nitrification on algal community and process performance

Complete nitrification was not positively or negatively associated with system performance (i.e., nutrient removal or algal biomass concentration), but imbalanced nitrifier communities resulted in partial nitrification and the accumulation of high levels of NO_2_^-^ (Figure 4A-B). The initial days of the first upset period (November 2021) were characterized by elevated influent NH_4_^+^ (> 80 mg N·L^-1^, Figure S2), acidic pH (5.5-6, grab sampling), and low alkalinity (< 100 mg·L^-1^ as CaCO_3_, Figure 1D), and the MRA of ammonia-oxidizing bacteria (AOB) was significantly (*p* < 0.05, ANOVA and Tukey’s HSD) higher (3.7-13.5%) than that of nitrite-oxidizing bacteria (NOB, 0.4-1%) (Figure 4A). At this high AOB:NOB ratio (>10), partial nitrification (conversion of NH_4_^+^ to NO_2_^-^) was followed by very little or no additional NO_3_^-^ formation. This led to the accumulation of NO_2_^-^ (> 40 mg N·L^-1^, grab sampling) which could pose a severe toxicity to the mixed microalgal community (Gonzalez-Camejo et al., 2020a). This nitrite accumulation along with other deleterious conditions could have been associated with the observed decline in biomass productivity and nutrient removal efficiency observed during upset period 1. To mitigate this issue, the alkalinity and SRT was increased (≥ 200 mg·L^-1^ as CaCO_3_ and 3-4 d, respectively) to aid in the recovery of NOBs as described in previous studies (Gonzalez-Camejo et al., 2020b).

**Figure 4.**
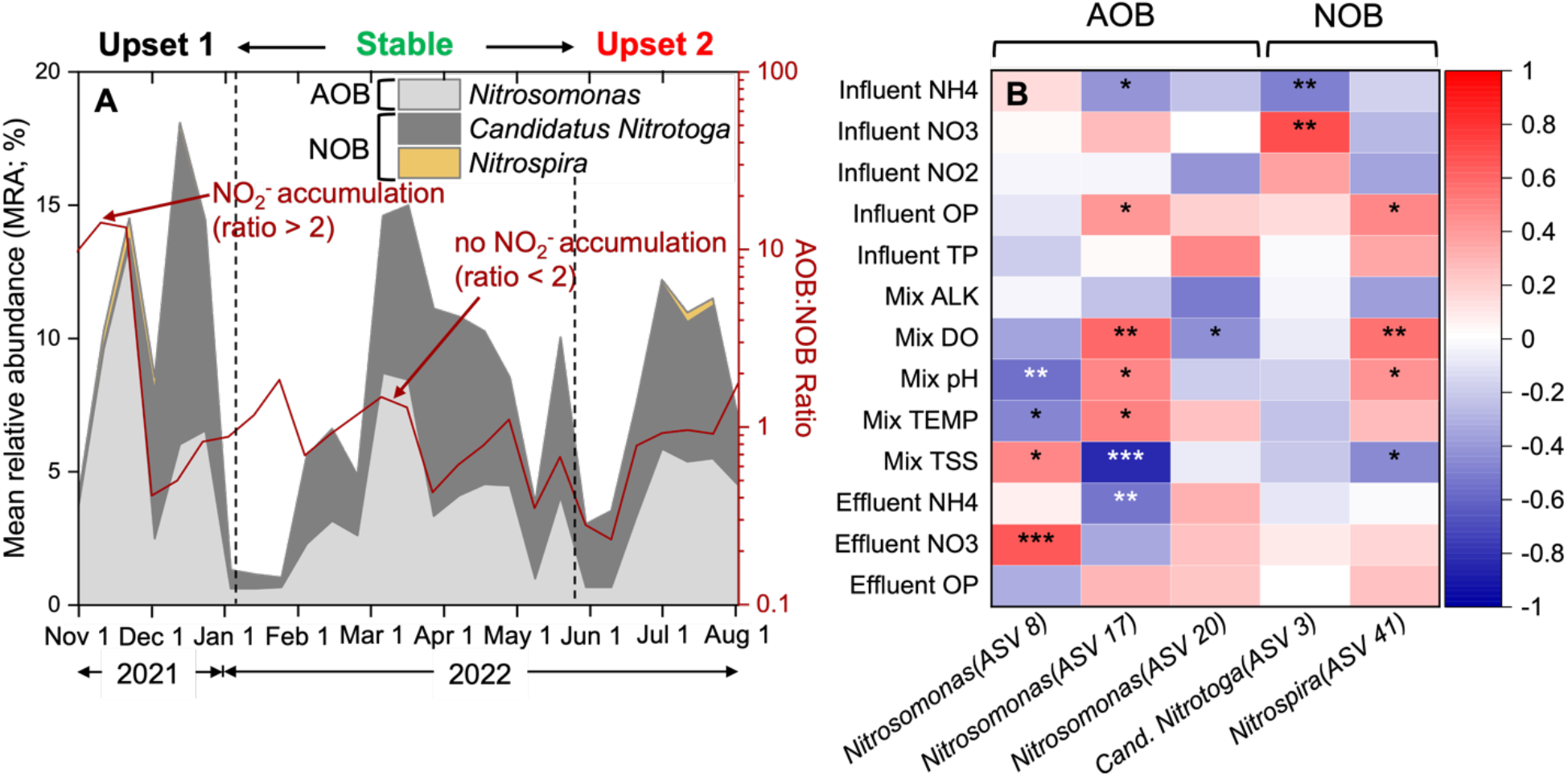
(A) Mean relative abundance (MRA) of ammonia and nitrite oxidizing bacteria (AOBs and NOBs) over time (left axis) and ratio of AOB and NOB over time (right axis). (B) Spearman (pairwise) correlation of the abiotic variables with specific AOB and NOB ASVs. Asterisks denote the *p-*value for each pair variable (no asterisk indicates *p* > 0.05, one asterisk indicates *p* < 0.05, two asterisks indicate *p* < 0.01, and three asterisks indicate *p* < 0.001. A balanced nitrifying community (AOB:NOB < 2) did not impact the system performance negatively, but imbalanced communities (AOB:NOB > 2) resulted in partial nitrification and nitrite levels that caused toxicity.

Under high ammonium concentrations and low alkalinity, as observed during the beginning of the first upset period, system pH dropped substantially (< 6) due to lack of alkalinity to counter the acidification due to ammonium oxidation (Faust et al., 2022). As pH decreases, a protonation of the NO_2_^-^ residual nitrite can lead to free nitrous acid (FNA) accumulation which can inhibit both AOBs and NOBs at low concentrations (Zhou et al., 2011). Although some acid tolerant AOBs can carry out ammonia oxidation at higher FNA concentrations (> 3 mg HNO_2_-N·L^-1^) (Wang et al., 2021), NOBs are very sensitive to low pH levels and can be completely inhibited by FNA at ppb levels (0.011 mg HNO_2_-N·L^-1^) (Vadivelu et al., 2006). In this study, the dominant AOB was *Nitrosomonas oligotropha*, which has a higher substrate affinity (*Km_(N. oligotropha)_* = 0.4-4.5 mg NH_4_-N·L^-1^, compared to the common *N. europaea*, *Km_(N. europaea)_* = 1.2-111 mg NH_4_-N·L^-1^) (Limpiyakorn et al., 2013; Sedlacek et al., 2019) and which has been shown to undergo physiological adaptations to carry out ammonia oxidation under more acidic conditions (pH = 4.5) than those that inhibit other AOB (*N. europaea,* pH < 6) (Fumasoli et al., 2017). In contrast, the presence of the *Nitrospira* NOBs were significantly (*p* < 0.05, method = Spearman) dependent on system pH and DO (Figure 4B), with low pH and DO limiting NO_2_^-^ oxidation and causing NO_2_^-^ accumulation. Long-term, a psychrotolerant NOB genus *Candidatus* Nitrotoga was selected over *Nitrospira*. However, this occurred after NO_2_^-^ and FNA achieved levels to pose severe toxicity to microalgae (> 25 mg NO_2_-N·L^-1^ (Aparicio et al., 2021) and >2 mg HNO_2_-N·L^-1^ FNA (Abbew et al., 2022)).

Mixed algal wastewater systems often try to limit nitrification (Krustok et al., 2016) to avoid NH_4_^+^ competition as a nutrient source for algal assimilation and as an electron donor for nitrifiers. While previous studies have shown that high nitrification levels (> 50% of the influent NH_4_^+^ concentration being converted to NO_3_^-^) can cause ammonium-limitation and reduce microalgal growth (González-Camejo et al., 2017), these conditions occur along with unfavorable microalgal growth conditions (e.g., high temperature (30-35 °C) and light intensity that causes photoinhibition (≥ 500 µmol·m^-2^·s^-1^) (Hasan Chowdury et al., 2020)). Indeed, the EcoRecover system experienced periods of high bacterial nitrification, with effluent NO_3_^-^ concentrations 35-70% of influent NH_4_^+^ during stable performance without adversely affecting microalgal growth and overall nutrient removal. In addition, while algae preferentially use NH_4_^+^ as a N source, they can also utilize converted NO_3_^-^ and organic N (Kumar and Bera, 2020). Here, the presence of AOB/NOB and nitrification had no significant correlation (positive or negative) with algal community structure and system performance, indicating that microalgae can co-exist with nitrifiers when provided with favorable growth conditions, including light (monthly average: 120-150 µmol·m^-2^·s^-1^) and alkalinity (monthly average: 280-315 mg·L^-1^ as CaCO_3_). While the presence or absence of nitrifiers did not significantly affect algal system performance, an imbalanced AOB/NOB community and resultant NO_2_^-^ accumulation was significantly associated deteriorating process performance and rapid shifts in algal community structure.

#### 3.3.2. Upset 2 - effect of zooplankton grazers and bacterial/fungal parasites on system performance

A range of potential pests were identified in the EcoRecover system (Table S2), which include zooplankton and other heterotrophic and mixotrophic grazers(Carney et al., 2016; Molina-Grima et al., 2022), fungal or fungal-like pathogens(Letcher et al., 2017a, 2017b), and bacterial pathogens (Lee et al., 2018), which have previously been reported to have negative effects on mixed microalgal systems. During upset period 2 (June 2022), a sudden shift in influent concentration (N:P ratio from > 25 to < 3) switched the EcoRecover system from P limited to N limited algal growth was associated with culture turnover from *Scenedesmus* to *Monoraphidium* (Figure 2A) which coincided with an increase in zooplankton grazers (e.g., ciliates and rotifers) (Figure 5A-B) and increase in ambient temperatures (from 13-17°C to 22-25 °C). A large increase in ciliates (0.4% to 11.8%) were observed (Figure 5B) as the system started to shift from *Scenedesmus* to *Monoraphidium* (Figure 2A). When the culture completely turned over to *Monoraphidium-* dominant (75%), the grazer communities exhibited a shift from ciliates to rotifers dominated by *Brachionus* sp. (9.9%). Subsequently, an increase in mixotrophic nano-flagellates dominated by *Ochromonas* sp. (2.6% to 10.1%) along with a decrease in *Monoraphidium* relative abundance (80.8% to 13.6%) and increase in relative abundance of Scenedesmus (0.2% to 13.7%), *Desmodesmus* (1.7% to 22.7%) and *Chlorella* (0.02% to 2.02%). The increase in *Ochromonas* also coincided with a decrease in cyanobacterial relative abundance on two separate occasions (2022-06-08 and 2022-07-13) (Figure 2B and 5B). As *Scenedesmus* relative abundance increased significantly (30.8%) towards the end of the upset period aided by operational interventions that increased the N:P ratio, a simultaneous increase in a fungal-like parasite *Aphelidium* (0.27% to 10.48%) was also observed. The abundance of pests dropped below the baseline (6.5%, representing the average total pest abundance observed during low pest prevalence, as shown in Figure 5A) by the end of the second upset period. The top zooplankton grazers or predators (≥ 1% MRA) included members from single-celled protozoa (e.g., ciliates, cercozoans) and multicellular animals (e.g., rotifers) including diverse ciliate genera (*Epistylis*, *Hemiophrys*, *Peritrichia*, and *Haptoria*), rotifers (*Brachionus)*, amoeba (*Rhogostoma*), and mixotrophic flagellates (*Ochromonas*, *Paraphysomonas*) (Figure 5C; Figure S5). Pathogens consisted of members from fungi or sister groups, including endoparasites *Aphelidium and LKM 11*, as well as bacterial parasites from the genera *Saprospira*, *Bdellovibrio*, *Oligoflexus*, *Vampirovibrio*, *Cytophaga*, *Bacillus*, and *Pseudomonas* (Figure S6).

**Figure 5.**
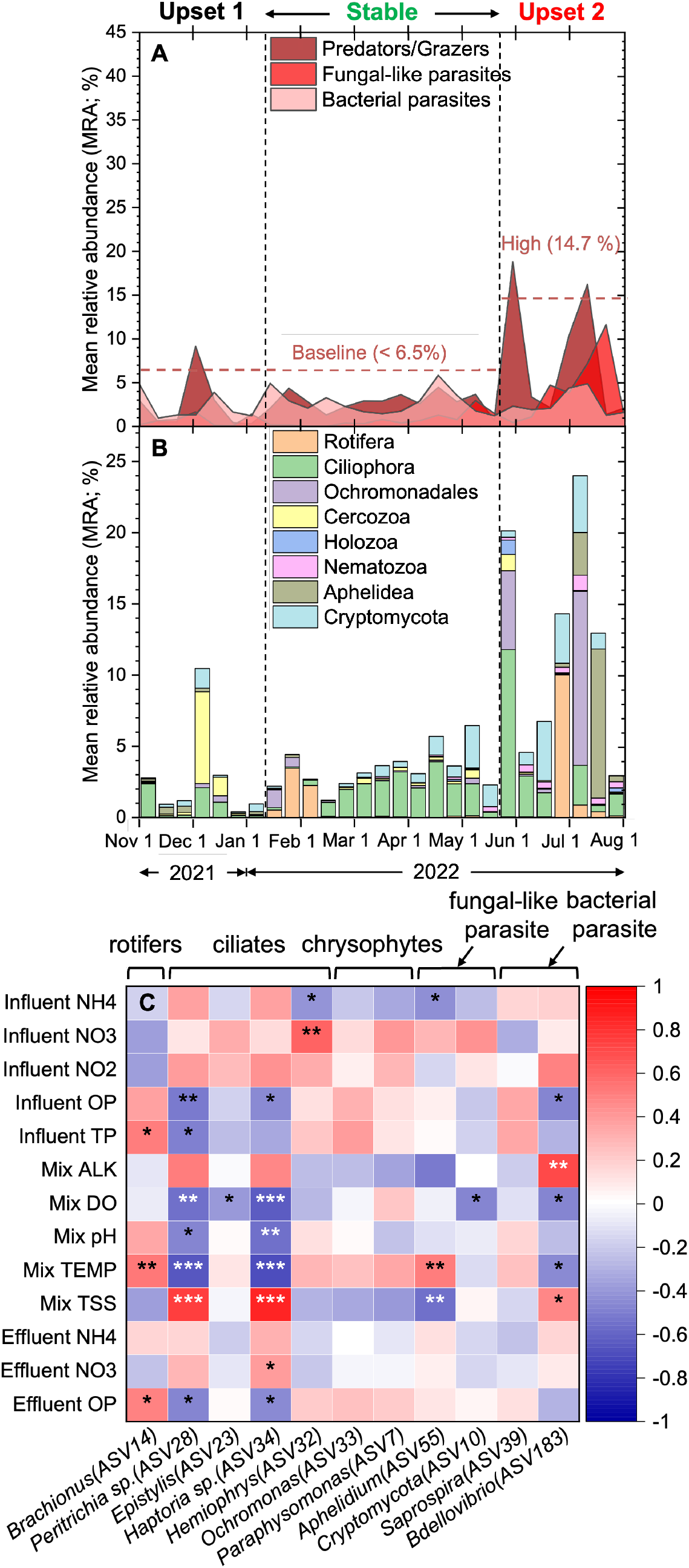
(A) Groups of potential pests including grazers or predators, fungal or fungal-like pathogens, and bacterial pathogens frequently observed in the EcoRecover system. (B) Predominant phyla of grazers or predators. (C) Spearman (pairwise) correlation of the abiotic variables with ASVs (≥ 1% mean relative abundance) representating potential pests. Asterisks denote the *p-*value for each pair variable (no asterisk indicates *p* > 0.05, one asterisk indicates *p* < 0.05, two asterisks indicate *p* < 0.01, and three asterisks indicate *p* < 0.001. Culture turnover from *Scenedesmus* to *Monoraphidium* was followed by high grazing activities of cilitates and rotifers during the second upset period that was characterized by nitrogen limiation (N:P < 3) and elevated temperature (22-25 °C). Grazer and pathogen dynamics showed specificity towards predominant taxa.

N:P ratio for *Scenedesmus* has been reported to range from 5-13:1, with ratios below this range demonstrating a reduction in both biomass productivity and nutrient removal efficiencies (Arbib et al., 2013; Xin et al., 2010). In contrast, *Monoraphidium* exhibits higher biomass productivity at lower N:P ratio in the range of 0.5-3:1 (Dhup and Dhawan, 2014) and has demonstrated maximum removal capacity even at low nitrogen concentrations [e.g., 0.1 mg N·L^-1^ (Khan et al., 2023)] such as those seen during the second upset period. Moreover, *Scenedesmus* may be more susceptible to N-limited stress than *Monoraphidium*, as the latter can produce unique biomolecules (e.g., tocopherols) to help maintain cell structure during periods of N-limitation (Singh et al., 2020). In any event, loss of *Scenedesmus* was coupled with subsequent grazing by ciliates that further reduced biomass. A complete turnover from *Scenedesmus* to *Monoraphidium* was coupled with grazer turnover from ciliates to rotifers (*Brachionus*), which could have resulted from the *Monoraphidium*-specific grazing activities of *Brachionus*, which have been shown to be ideal for their feeding due to the equivalent spherical diameter (ESD) of *Monoraphidium* sp. (1.5-6.5 µm) (Rothhaupt, 1990). Contrarily, *Brachionus* could also have switched from feeding on *Scenedesmus* to ciliates due to the scarcity of microalgal cells and available nutrients (Gilbert and Jack, 1993). As P limitation was reinstated, rotifers were replaced by the mixotrophic algal nano-flagellate *Ochromonas.* Subsequently, as *Scenedesmus* recovered in the system, a fungal or fungal-like endobiotic pathogen, *Aphelidium*, quickly became prevalent, consistent with its known specific parasitism on *Scenedesmus* as described in previous studies (Letcher et al., 2017a, 2017c). Of all the pests, at least two species (*Brachionus* and *Aphelidium*) showed positive correlation (*p* ≤ 0.01, method = Spearman) with the temperature of the mixed system (Figure 5C) and elevated summer temperatures (22-25 °C) (Ferro et al., 2020). In general, these pests persisted within the system across seasons, even during stable performance periods, but their abundance was not significantly associated with changes in process performance or community structure (baseline abundance < 6.5%), and only exhibited increased grazing or pathogenic activities during favorable growth conditions characterized by elevated summer temperature and microalgal stress. Overall, the dynamics of the zooplankton and mixotrophic grazers and fungal or fungal-like parasites observed in this study indicated their specificity towards specific taxa in the system (Figure 5B). To minimize culture upset events, future studies should adopt a comprehensive strategy for pest control, involving an efficient and rapid detection of pests and their associated algal phenotypic changes.

## 4. Conclusion

Overall, this study highlights the importance of key operational and environmental factors at the full-scale for mixed microalgal nutrient recovery. In keeping with our hypothesis, high-throughput sequencing of the 18S and 16S rRNA genes revealed distinct eukaryotic and bacterial community dynamics across periods of system upset and stable performance and identified associated abiotic drivers during these time periods. Key takeaways include:

- Stable operational periods were characterized by low eukaryotic diversity (*D_INVSIMPSON_* = 2.01) and a dominant *Scenedesmus* population (55-80%), while upset periods had higher algal diversity (*D_INVSIMPSON_* = 3.04 and 4.5 during upset 1 and upset, respectively) and were dominated by *Chlorella* and *Monoraphidium*.
- Long-term cultures contained highly diverse bacterial communities across both upset and stable periods, including nitrifiers (*D_INVSIMPSON_* = 19.1), but Cyanobacteria dominated during stable performance (30-56%).
- A balanced nitrifying community did not positively or negatively impact microalgae, but partial nitrification resulting in NO_2_^-^ accumulation, insufficient alkalinity (< 100 mg·L^-1^ as CaCO_3_), and low pH (< 6) were associated with deteriorating process performance and culture turnover.
- The stable algal community was positively correlated (*p* < 0.05, method = Spearman) with several key abiotic factors, including system pH, alkalinity, and influent ammonium (NH_4_^+^) which promoted increased algal biomass and nutrient assimilation.
- Pest temporal dynamics followed host specific dynamics and occurred during elevated summer temperature (22-25 °C) after the loss of system performance and community turnover, revealing the importance of maintaining healthy algal cultures and high biomass productivity.

## Acknowledgements

This work has received support from the U.S. Department of Energy, Office of Energy Efficiency and Renewable Energy, under Award Number DE-EE0009270, and the authors express gratitude to Jeremy Guest and Hannah Molitor from UIUC for project management and overall team support, as well as Autumn Fisher and Kevin McGraw from Clearas Water Recovery for their substantial contributions to operating and maintaining the EcoRecover system. Special appreciation goes to John Bond, Director of Public Works at the Village of Roberts, and the Public Works staff for their invaluable on-site support and expertise.

