## Supporting Information for "Community Structure and Function During Periods of High Performance and System Upset in a Full-Scale Mixed Microalgal Wastewater Resource Recovery Facility"

**List of Figures**

Figure S1 (p. 2)
Figure S2 (p. 2)
Figure S3 (p. 3)
Figure S4 (p. 4)
Figure S5 (p. 5)
Figure S6 (p. 6)
Figure S7 (p. 7)
Figure S8 (p. 8)
Figure S9 (p. 9)

**List of Tables**

Table S1 (p. 10)
Table S2 (p. 11-12)
Table S3 (p. 13-14)
Table S4 (p. 15-16)

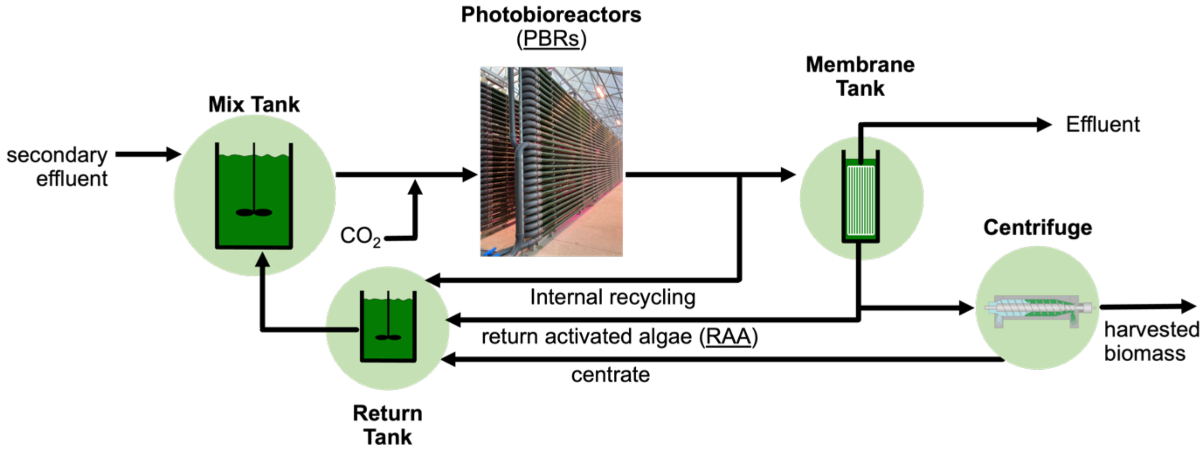


**Figure S1:** Schematic of the major operational units in the full-scale EcoRecover system.

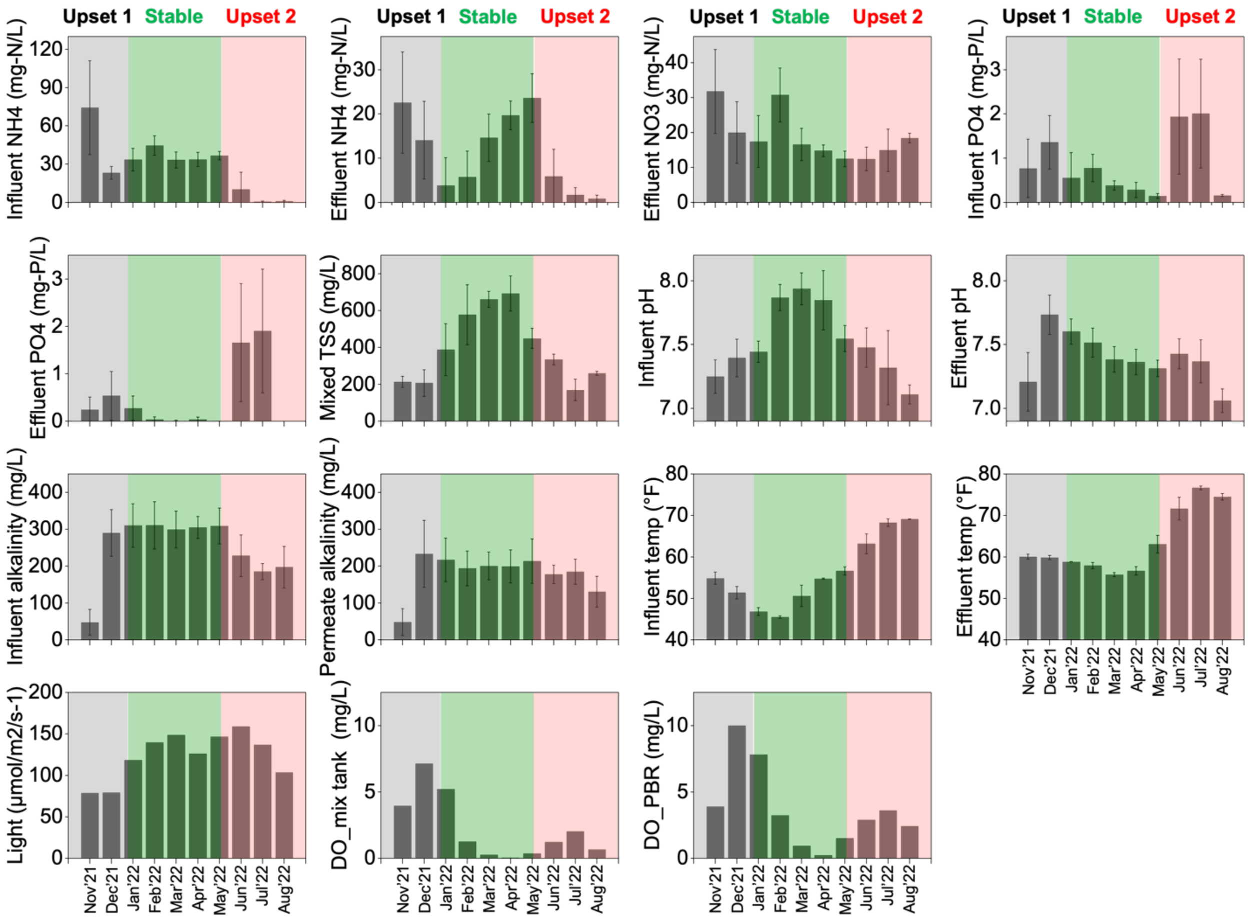


**Figure S2:** Average monthly operational parameters of the EcoRecover system during the long-term monitoring period (Nov 2021-Aug 2022).

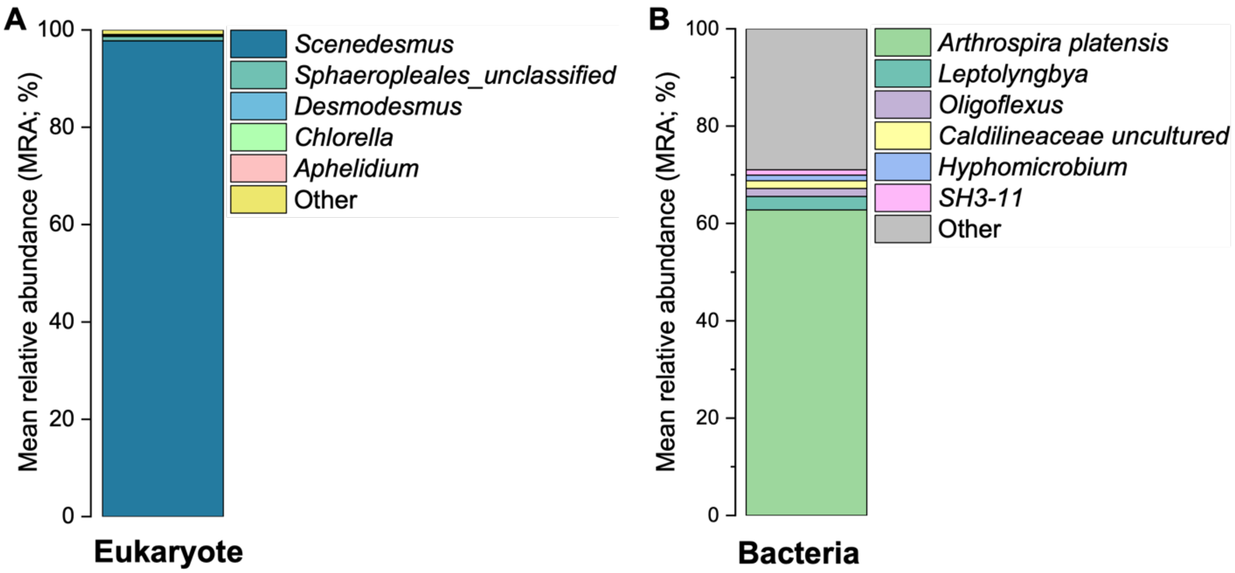


**Figure S3:** Mean relative abundance (MRA) of the predominant (A) eukaryotic and (B) bacterial genera present in the algae paste sourced from an EcoRecover system located in the South Davis Sewer District (SDSD) in Utah. The algae paste was inoculated in the EcoRecover system at Village of Roberts, WI, in early December 2021, alongside significant operational changes (detailed in **section 3.3.1**), which contributed to the recovery of the system from culture crash.

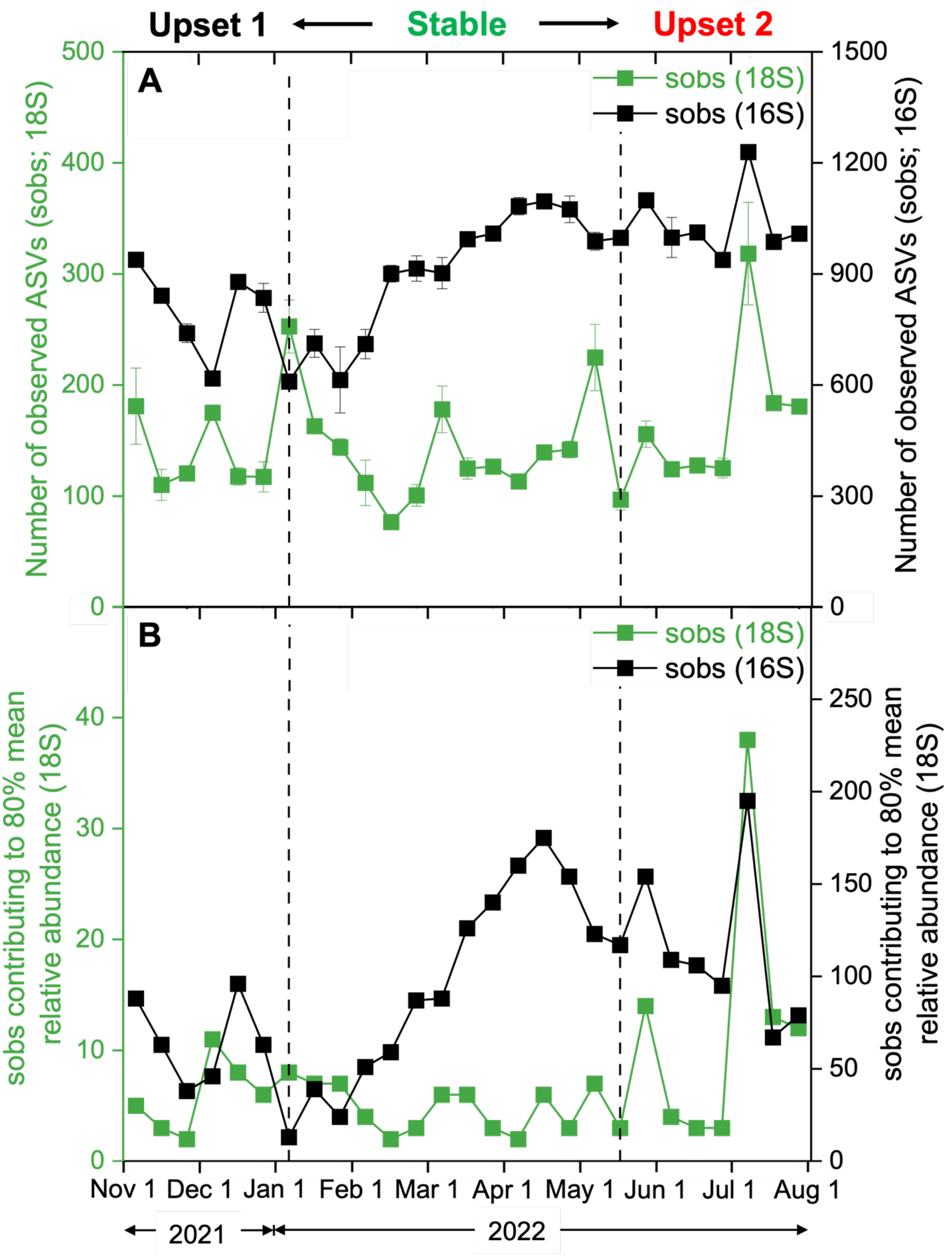

Upset 1
Stable
Upset 2
A
sobs (18S)
sobs (16S)
Number of observed ASVs (sobs; 18S)
Number of observed ASVs (sobs; 16S)
B
sobs contributing to 80% mean relative abundance (18S)
Nov 1
Dec 1
Jan 1
Feb 1
Mar 1
Apr 1
May 1
Jun 1
Jul 1
Aug 1
2021
2022

**Figure S4:** Number of observed ASVs (Sobs) identified from the 18S rRNA (left axis, green) and 16S rRNA (right axis, black) gene sequencing contributing to (A) the total mean relative abundance (MRA) of each sample and (B) 80% of the MRA across the periods of stable performance and system upsets. Bacterial communities maintained 5 – 7 times higher Sobs than eukaryotes. The average Sobs of stable eukaryotic community were relatively lower than communities during upset events [$D_{sobs}$ *(avg)* = 134, 153, 174 during stable, upset 1, and upset 2 periods, respectively. For bacterial community, Sobs contributing to 80% MRA was significantly less ($p < 0.05$, ANOVA and Tukey's HSD) during upset 1 compared to stable and upset 2 periods [$D_{sobs}$ *(avg)* = 58, 103, 116 during upset 1, stable, and upset 2, respectively).

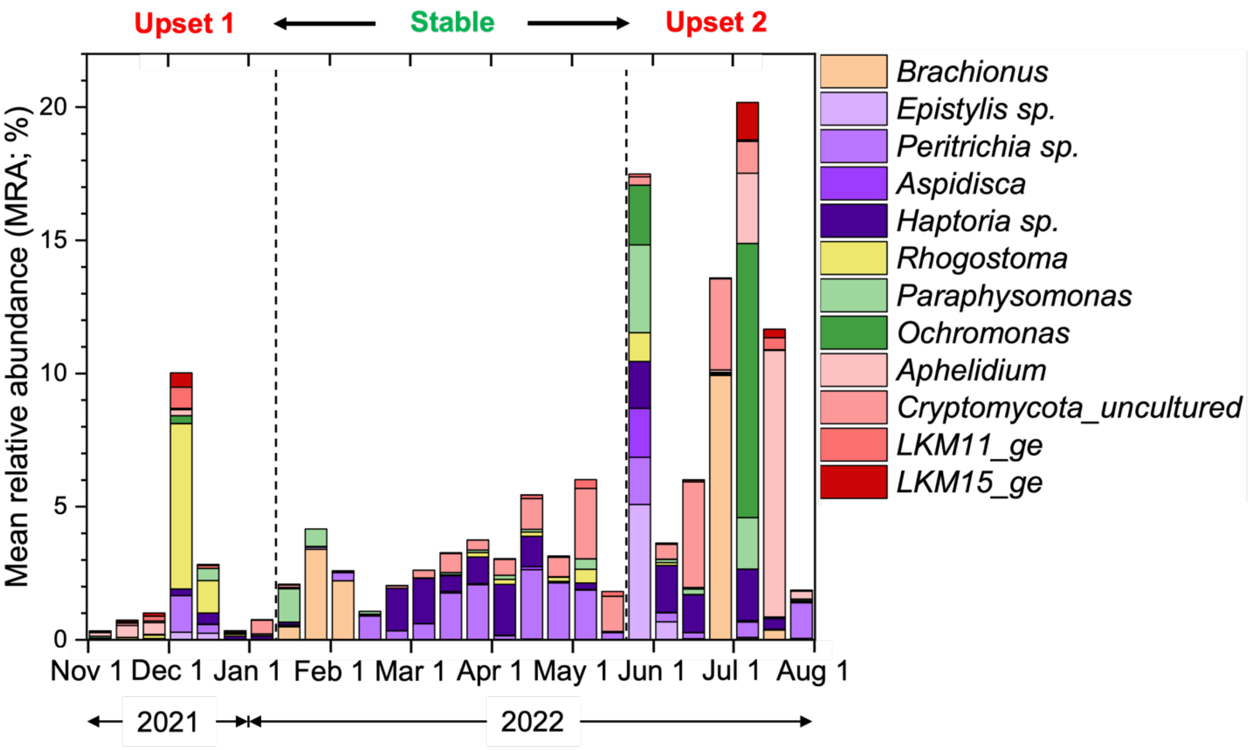


**Figure S5:** Mean relative abundance (MRA) of the top (≥ 1%) pest genera identified from 18S rRNA gene sequencing, including zooplankton grazers, mixotrophic predators and fungal or fungal-like pathogens. High grazing activities dominated by ciliates, rotifers, flagellates, and fungal-like pathogen were observed right after the culture turnover from *Scenedesmus* to *Monoraphidium* during the second upset period. The dynamics of these grazers and pathogens showed specificity towards predominant taxa present in the system.

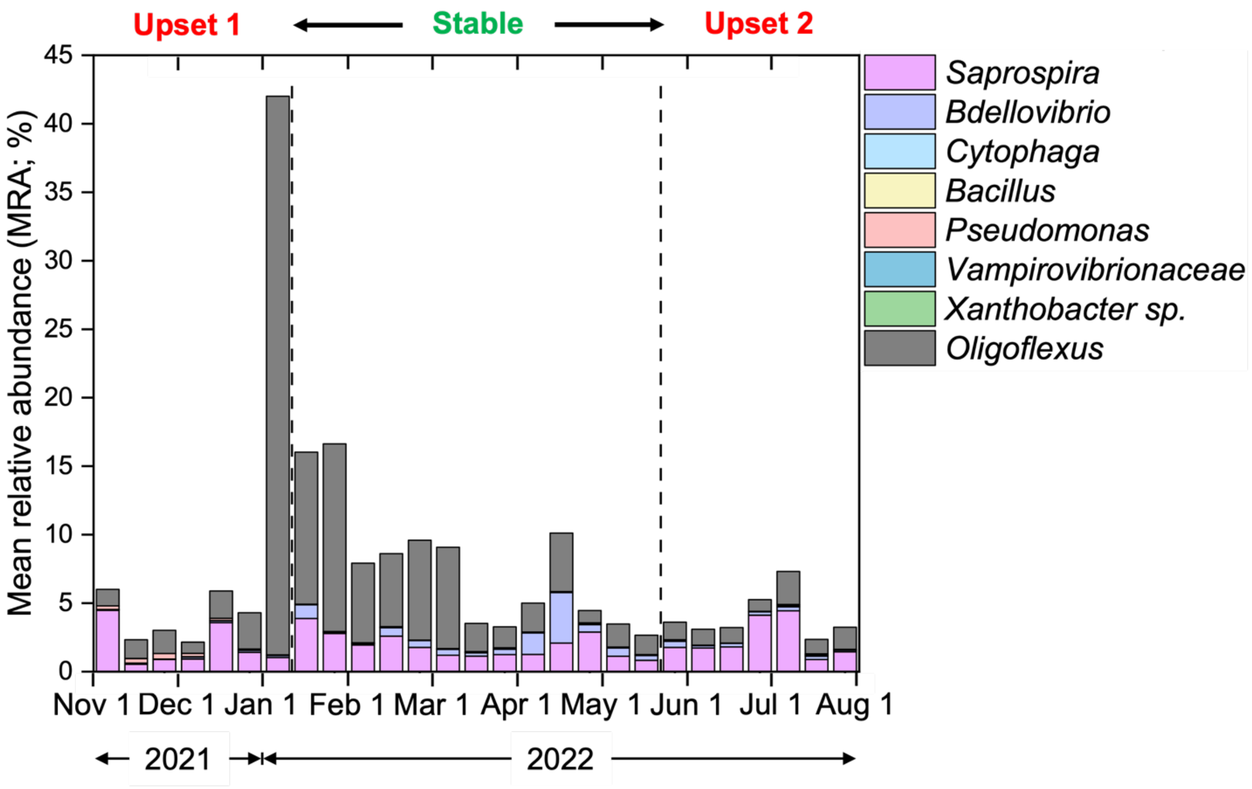


**Figure S6:** Mean relative abundance (MRA) of the top pest genera (≥ 1%) identified from 16S rRNA gene sequencing. All potential bacterial pests remained below the baseline abundance (6.5%) across sampling periods, except for the genus *Oligoflexus*, a member of the proteobacterial class *Oligoflexia* and order *Bdellovibrionales*, which includes *Bdellovibrio*-like predatory bacteria (Waite et al., 2020). It should be noted that, *Oligoflexus* was not included in the bacterial pest category in the main text (Figure 4A), since it was not associated with loss of microalgal biomass in this study. Although the genus *Oligoflexus* hasn't been confirmed as a microalgae specific pest yet, a recent study reported an uncultured *Bdellovibrio*-like bacterial predator likely from the same order (*Bdellovibrionales*) causing the culture loss of *Nannochloropsis* in outdoor ponds (Lee et al., 2018).

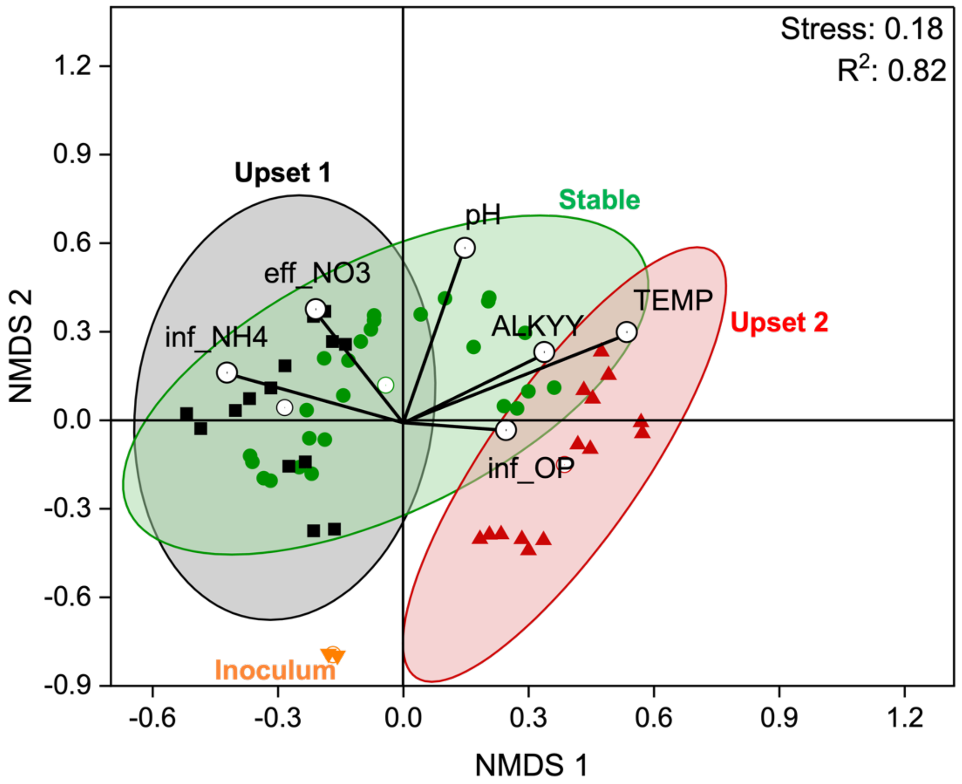


**Figure S7:** Non-metric multi-dimensional scaling (NMDS) plot using Bray-Curtis dissimilarity showing distances among bacterial communities during periods of stable performance and upset events, with lines indicating strength and direction of correlation (method = Spearman) for each variable. Ellipses containing 95% of cluster-assigned data points show greater overlap between stable and upset 1 communities compared to upset 2. The stable community was associated with pH, alkalinity, influent $NH_4^+$, and effluent $NO_3^-$.

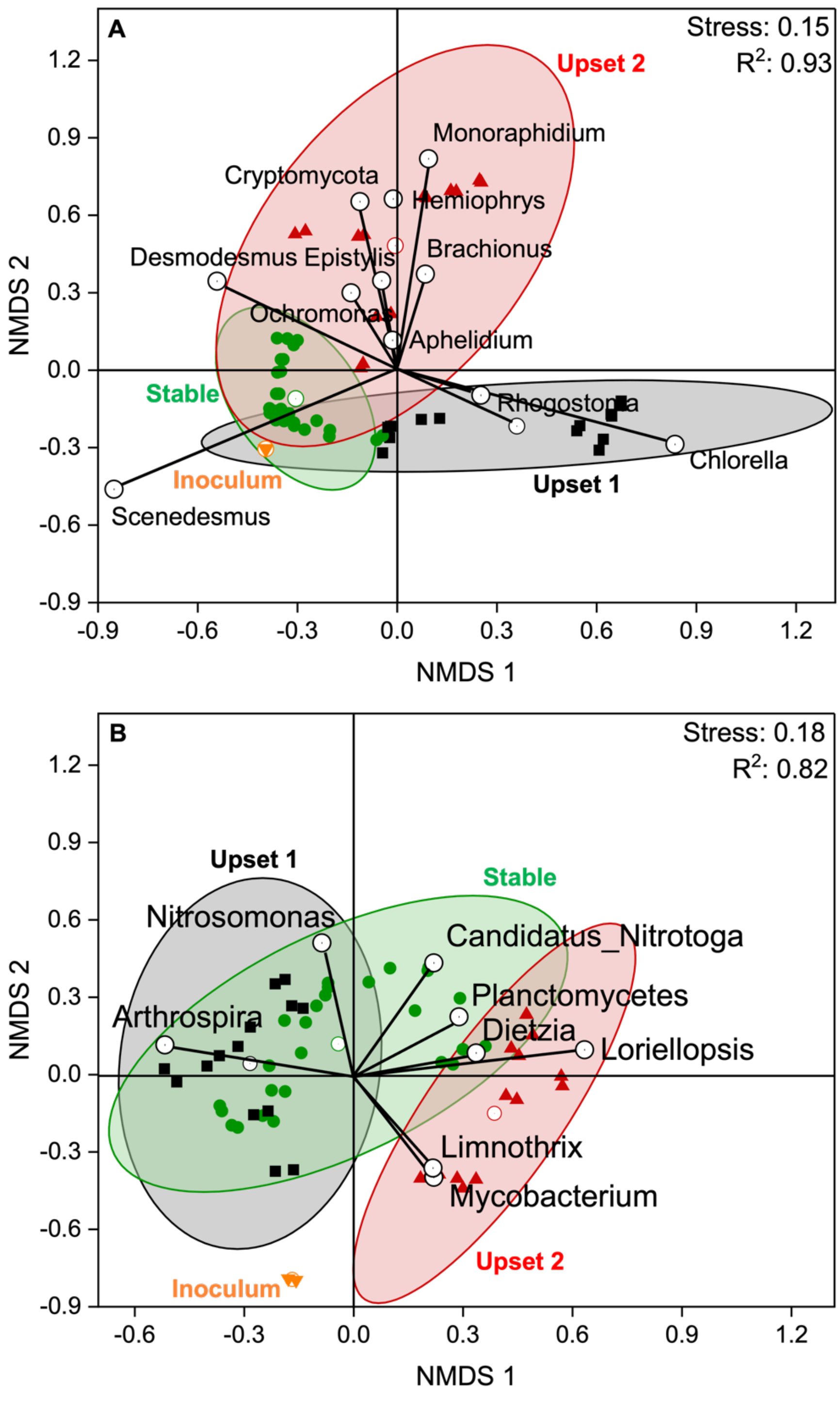


**Figure S8:** Non-metric multi-dimensional (NMDS) plot using Bray-Curtis dissimilarity showing distances among (A) eukaryotic and (B) bacterial communities, with lines indicating strength and direction of correlation (method = Spearman) of top ASVs that contributed to community turnovers during periods of stable performance and upset events.

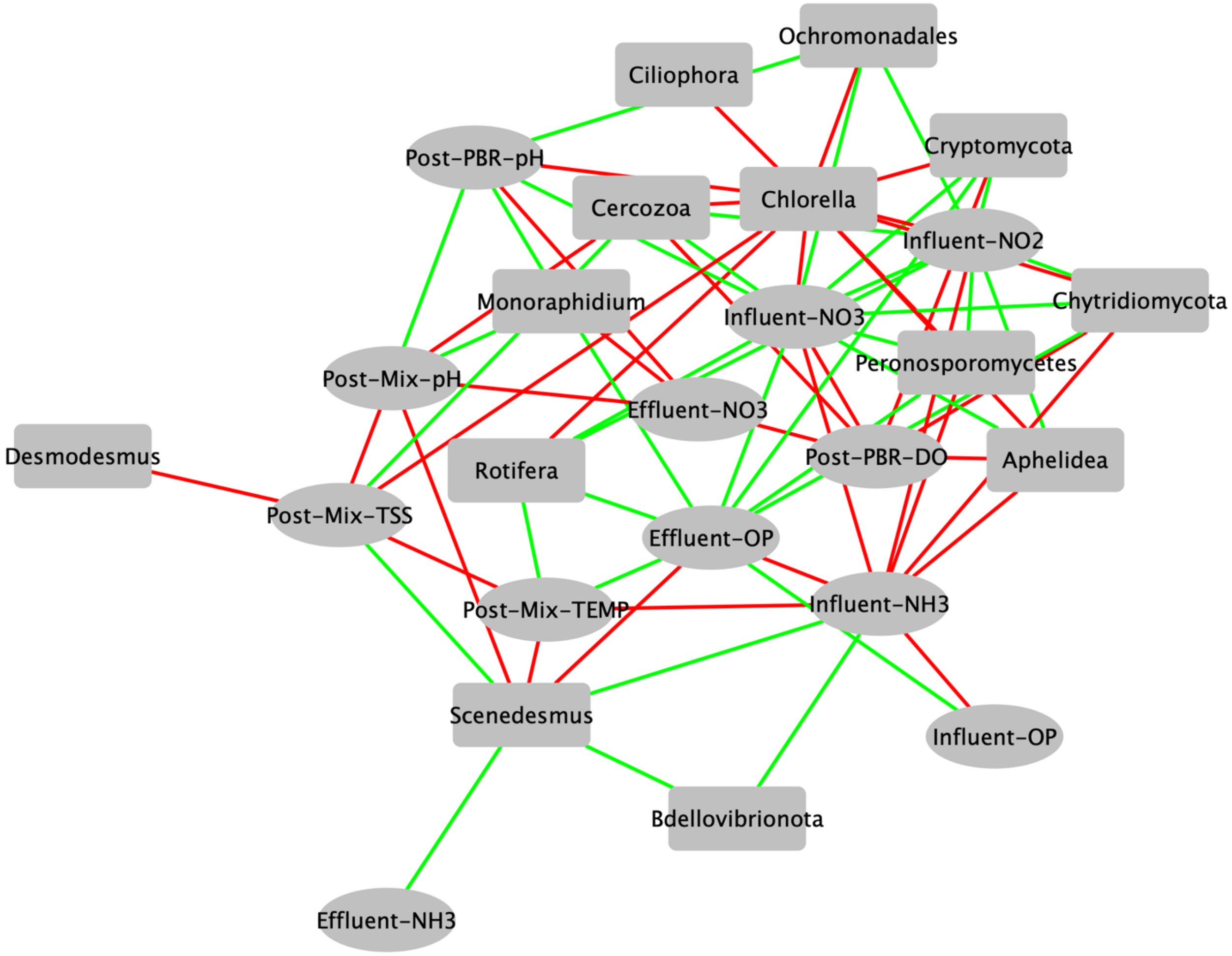


**Figure S9:** Co-occurrence network (Co-Net) analysis of 18S community structure and abiotic functional variables. The network consists of eukaryotic taxa that strongly ($p < 0.05$, Pearson) correlate with the metadata. The direction of interactions is color coded [positive (green) and negative (red)].

**Table S1.** On-site laboratory measurements performed on grab samples.

| **Measurement [units]** | **Hach kits (measuring range)** |
|---|---|
| Alkalinity [$mg \cdot L^{-1}$ $CaCO_3$] | TNT870 (25-400), |
| Ammonium [$mg\text{-}N \cdot L^{-1}$] | TNT830 (0.015-2), TNT831 (1-12), and TNT832 (2-47) |
| Nitrate [$mg\text{-}N \cdot L^{-1}$] | TNT835 (0.23-13.5) and TNT836 (5-35) |
| Nitrite [$mg\text{-}N \cdot L^{-1}$] | TNT839 (0.015-0.6) and TNT840 (0.6-6) |
| Total nitrogen [$mg\text{-}N \cdot L^{-1}$] | TNT827 (5-40) |
| Orthophosphate [$mg\text{-}P \cdot L^{-1}$] | TNT843 (0.05-1.5) |
| Total phosphate [$mg\text{-}P \cdot L^{-1}$] | TNT843 (0.05-1.5) |

**Table S2:** Taxonomic information for the ASVs representing harmful pests.

| Category | Group | Phylum | ASV # | Taxonomy | | | | | |
|---|---|---|---|---|---|---|---|---|---|
| Predator/Grazer | Zooplanknton (Animalia) | Rotifer | ASV 14 | Eukaryota(100) | Rotifera(100) | Monogononta(100) | Ploimida(100) | Ploimida_fa(100) | Brachionus(100) |
| | | | ASV 70 | Eukaryota(100) | Rotifera(100) | Monogononta(100) | Ploimida(100) | Ploimida_fa(100) | Ploimida_ge(100) |
| | | | ASV 87 | Eukaryota(100) | Rotifera(100) | Bdelloidea(100) | Adinetida(100) | Adinetida_fa(100) | Adinetida_ge(100) |
| | | | ASV 161 | Eukaryota(100) | Rotifera(100) | Monogononta(100) | Monogononta _unclass(100) | Monogononta _unclass(100) | Monogononta_ unclass(100) |
| | | Nematozoa | ASV 67 | Eukaryota(100) | Nematozoa(100) | Chromadorea(100) | Rhabditida(100) | Rhabditida_fa(100) | Rhabditida_ge(100) |
| | | | ASV 325 | Eukaryota(100) | Nematozoa(100) | Chromadorea(100) | Araeolaimida(100) | Araeolaimida_fa(100) | Araeolaimida_ ge(100) |
| | Zooplanknton (Protozoa) | Ciliophora | ASV 23 | Eukaryota(100) | Ciliophora(100) | Intramacronucleata(100) | Conthreep(100) | Oligohymenophorea(100) | Oligohymenophorea _unclass(100) |
| | | | ASV 28 | Eukaryota(100) | Ciliophora(100) | Intramacronucleata(100) | Conthreep(100) | Oligohymenophorea(100) | Peritrichia_ge(100) |
| | | | ASV 32 | Eukaryota(100) | Ciliophora(100) | Intramacronucleata(100) | Litostomatea(100) | Haptoria(100) | Hemiophrys(100) |
| | | | ASV 34 | Eukaryota(100) | Ciliophora(100) | Intramacronucleata(100) | Litostomatea(100) | Haptoria(100) | Haptoria_ unclass(100) |
| | | | ASV 38 | Eukaryota(100) | Ciliophora(100) | Intramacronucleata(100) | Litostomatea(100) | Haptoria(100) | Loxophyllum(100) |
| | | | ASV 52 | Eukaryota(100) | Ciliophora(100) | Intramacronucleata(100) | Spirotrichea(100) | Euplotia(100) | Aspidisca(100) |
| | | | ASV 62 | Eukaryota(100) | Ciliophora(100) | Intramacronucleata(100) | Conthreep(100) | Oligohymenophorea(100) | Dexiotricha(100) |
| | | | ASV 77 | Eukaryota(100) | Ciliophora(100) | Intramacronucleata(100) | Conthreep(100) | Oligohymenophorea(100) | Telotrochidium(100) |
| | | | ASV 85 | Eukaryota(100) | Ciliophora(100) | Intramacronucleata(100) | Litostomatea(100) | Haptoria(100) | Amphileptus(100) |
| | | Cercozoa | ASV 25 | Eukaryota(100) | Cercozoa(100) | Thecofilosea(100) | Cryomonadida(100) | Rhizaspididae(100) | Rhogostoma(100) |
| | | | ASV 125 | Eukaryota(100) | Cercozoa(100) | Cercozoa_unclass(100) | Cercozoa _unclass(100) | Cercozoa_unclassified(100 ) | Cercozoa_ unclass(100) |
| | Algal flagellate | Ochrophyta | ASV 7 | Eukaryota(100) | Ochrophyta(100) | Chrysophyceae(100) | Ochromona -dales(100) | Ochromonadales_fa(100) | Paraphysomonas(100 ) |
| | | | ASV 33 | Eukaryota(100) | Ochrophyta(100) | Chrysophyceae(100) | Ochromona -dales(100) | Ochromonadales_fa(100) | Ochromonas(100) |
| | | | ASV 396 | Eukaryota(100) | Ochrophyta(100) | Chrysophyceae(100) | Chromulinales(100) | Chromulinales_fa(100) | Poterioochro- monas(100) |
| | | Holozoa | ASV 71 | Eukaryota(100) | Holozoa(100) | Choanoflagellida(100) | Craspedida_or(100) | Salpingoecidae(100) | Salpingoeca(100) |
| | | | ASV 183 | Eukaryota(100) | Holozoa(100) | Choanoflagellida(100) | Choanoflagellida _unclass(100) | Choanoflagellida _unclass(100) | Choanoflagellida _unclass(100) |
| | | | ASV 202 | Eukaryota(100) | Holozoa(100) | Choanoflagellida(100) | Codonosigidae(100) | Codonosigidae_fa(100) | Monosiga(100) |

| | | | | | | | | | |
|---|---|---|---|---|---|---|---|---|---|
| | | | ASV 307 | Eukaryota(100) | Holozoa(100) | Choanoflagellida(100) | Craspedida_or(100) | Craspedida _unclass(100) | Craspedida _unclass(100) |
| | | Protalveo -lata | ASV 430 | Eukaryota(100) | Protalveolata(100) | Colpodellida(100) | Colpodellida_or(100) | Colpodellida_fa(100) | Colpodella(100) |
| Pathogen/Parasite | Fungi or sister group | Cryptomy -cota | ASV 9 | Eukaryota(100) | Eukaryota _unclass(100) | Eukaryota_unclass(100) | Eukaryota _unclass(100) | Eukaryota _unclass(100) | Cryptomycota _unclass(100) |
| | | | ASV 10 | Eukaryota(100) | Eukaryota _unclass(100) | Eukaryota_unclass(100) | Eukaryota _unclass(100) | Eukaryota _unclass(100) | Cryptomycota _unclass(100) |
| | | | ASV 86 | Eukaryota(100) | Cryptomycota(100) | LKM11_cl(100) | LKM11_or(100) | LKM11_fa(100) | LKM11_ge(100) |
| | | | ASV 102 | Eukaryota(100) | LKM15(100) | LKM15_cl(100) | LKM15_or(100) | LKM15_fa(100) | LKM15_ge(100) |
| | | Aphelidea | ASV 16 | Eukaryota(100) | Aphelidea(100) | Aphelidea_cl(100) | Aphelidea_or(100) | Aphelidea_fa(100) | uncultured(100) |
| | | | ASV 36 | Eukaryota(100) | Aphelidea(100) | Aphelidea_cl(100) | Aphelidea_or(100) | Aphelidea_fa(100) | Aphelidea _unclass(100) |
| | | Chytridio -mycota | ASV 65 | Eukaryota(100) | Chytridio -mycota(100) | Chytridiomycetes(100) | Chytridiomycetes _unclass(100) | Chytridiomycetes _unclass(100) | Chytridiomycetes _unclass(100) |
| | | Peronospor o-mycetes | ASV 13 | Eukaryota(100) | Peronosporomy -cetes(100) | Peronosporomy -cetes_cl(100) | Peronosporo- mycetes_or(100) | Peronosporo -mycetes(100) | Paralagenidium(99) |
| | | | ASV 113 | Eukaryota(100) | Peronosporomy- cetes(100) | Peronosporomy -cetes_cl(100) | Peronosporo- mycetes_or(100) | Peronosporo -mycetes(100) | Peronosporomycetes _unclass(100) |
| | Bacteria | Bdellovibri -onota | ASV 6 | Bacteria(100) | Bdellovibrio -nota(100) | Oligoflexia(100) | Oligoflexales(100) | Oligoflexaceae(100) | Oligoflexus(100) |
| | | | ASV 68 | Bacteria(100) | Bdellovibrio -nota(100) | Bdellovibrionia(100) | Bdellovibrio -nales(100) | Bdellovibrionaceae(100) | OM27_clade(100) |
| | | | ASV 183 | Bacteria(100) | Bdellovibrio -nota(100) | Bdellovibrionia(100) | Bdellovibrio -nales(100) | Bdellovibrionaceae(100) | Bdellovibrio(100) |
| | | | ASV 237 | Bacteria(100) | Bdellovibrio -nota(100) | Oligoflexia(100) | Oligoflexales(100) | Oligoflexaceae(100) | uncultured(100) |
| | | | ASV 363 | Bacteria(100) | Bdellovibrio -nota(100) | Oligoflexia(100) | Oligoflexales(100) | Oligoflexaceae(100) | Oligoflexus(100) |
| | | | ASV 412 | Bacteria(100) | Bdellovibrio -nota(100) | Oligoflexia(100) | 053A03-B-DI -P58(100) | 053A03-B-DI-P58(100) | 053A03-B-DI- P58_ge(100) |
| | | Cyanobac -teria | ASV 3459 | Bacteria(100) | Cyanobac -teria(100) | Vampirivibrionia(100) | Vampirovibrio- nales(100) | Vampirovibrionales _unclass(100) | Vampirovibrionales _unclass(100) |
| | | | ASV 6423 | Bacteria(100) | Cyanobac -teria(100) | Vampirivibrionia(100) | Vampirovibrio- nales(100) | Vampirovibrio- naceae(100) | Vampirovibrio(100) |
| | | | ASV 13351 | Bacteria(100) | Cyanobac -teria(100) | Vampirivibrionia(100) | Vampirovibrio- nales(100) | Vampirovibrio- naceae(100) | Vampirovibriona -ceae_ge(100) |
| | | Bacteroi -dota | ASV 1869 | Bacteria(100) | Bacteroi -dota(100) | Bacteroidia(100) | Cytophagales(100) | Cytophagaceae(100) | Cytophaga(100) |

**Table S3:** Taxonomic information for the top 50 ASVs from 18S rRNA gene sequencing.

| ASV | Size | Taxonomy | | | | | |
|---|---|---|---|---|---|---|---|
| ASV 1 | 438261 | Eukaryota(100) | Chlorophyta_ph(100) | Chlorophyceae(100) | Sphaeropleales(100) | Sphaeropleales_fa(100) | Scenedesmus(100) |
| ASV 2 | 139745 | Eukaryota(100) | Chlorophyta_ph(100) | Trebouxiophyceae(100) | Chlorellales(100) | Chlorellales_fa(100) | Chlorella(100) |
| ASV 3 | 129699 | Eukaryota(100) | Chlorophyta_ph(100) | Chlorophyceae(100) | Sphaeropleales(100) | Sphaeropleales_fa(100) | Monoraphidium(100) |
| ASV 4 | 96241 | Eukaryota(100) | Chlorophyta_ph(100) | Chlorophyceae(100) | Sphaeropleales(100) | Sphaeropleales_fa(100) | Desmodesmus(100) |
| ASV 5 | 65429 | Eukaryota(100) | Chlorophyta_ph(100) | Chlorophyceae(100) | Sphaeropleales(100) | Sphaeropleales_fa(100) | Desmodesmus(100) |
| ASV 6 | 11807 | Eukaryota(100) | Chlorophyta_ph(100) | Chlorophyceae(100) | Sphaeropleales(100) | Sphaeropleales_fa(100) | Kirchneriella(99) |
| ASV 7 | 10818 | Eukaryota(100) | Ochrophyta_ph(100) | Chrysophyceae(100) | Ochromonadales(100) | Ochromonadales_fa(100) | Paraphysomonas(100) |
| ASV 8 | 10314 | Eukaryota(100) | Chlorophyta_ph(100) | Trebouxiophyceae(100) | Chlorellales(100) | Chlorellales_fa(100) | Chlorella(100) |
| ASV 9 | 8905 | Eukaryota(100) | Eukaryota_unclass.(100) | Eukaryota_unclass.(100) | Eukaryota_unclass.(100) | Eukaryota_unclass.(100) | Eukaryota_unclass.(100) |
| ASV 10 | 6776 | Eukaryota(100) | Eukaryota_ unclass.(100) | Eukaryota_unclass.(100) | Eukaryota_unclass.(100) | Eukaryota_unclass.100) | Cryptomycota(98) |
| ASV 11 | 5829 | Eukaryota(100) | Chlorophyta_ph(100) | Chlorophyceae(100) | Sphaeropleales(100) | Sphaeropleales_fa(100) | Coelastrella(100) |
| ASV 12 | 5734 | Eukaryota(100) | Chlorophyta_ph(100) | Chlorophyceae(100) | Sphaeropleales(100) | Sphaeropleales_fa(100) | Desmodesmus(100) |
| ASV 13 | 5592 | Eukaryota(100) | Peronosporomycetes(100) | Peronosporomycetes(100) | Peronosporomycetes(100) | Peronosporomycetes_fa(100) | Paralagenidium(99) |
| ASV 14 | 5207 | Eukaryota(100) | Rotifera(100) | Monogononta(100) | Ploimida(100) | Ploimida_fa(100) | Brachionus(100) |
| ASV 15 | 4856 | Eukaryota(100) | Cercozoa(100) | Cercozoa_unclass.(100) | Cercozoa_unclass.(100) | Cercozoa_unclass.(100) | Lecythium(96) |
| ASV 16 | 3917 | Eukaryota(100) | Aphelidea(100) | Aphelidea_cl(100) | Aphelidea(100) | Aphelidea_fa(100) | uncultured(100) |
| ASV 17 | 3688 | Eukaryota(100) | Eukaryota_ unclass.(100) | Eukaryota_unclass.(100) | Eukaryota_unclass.(100) | Eukaryota_unclass.(100) | Eukaryota_unclass.(100) |
| ASV 18 | 3353 | Eukaryota(100) | Diatomea(100) | Bacillariophyceae(100) | Bacillariophyceae(100) | Bacillariophyceae_fa(100) | Nitzschia(100) |
| ASV 19 | 3097 | Eukaryota(100) | Chlorophyta_ph(100) | Trebouxiophyceae(100) | Chlorellales(100) | Chlorellales_fa(100) | Chlorella(100) |
| ASV 20 | 2963 | Eukaryota(100) | Chlorophyta_ph(100) | Chlorophyceae(100) | Sphaeropleales(100) | Sphaeropleales_fa(100) | Desmodesmus(100) |
| ASV 21 | 2755 | Eukaryota(100) | Diatomea(100) | Bacillariophyceae(100) | Bacillariophyceae(100) | Bacillariophyceae_fa(100) | Gomphonema(100) |
| ASV 22 | 2528 | Eukaryota(100) | Chlorophyta_ph(100) | Trebouxiophyceae(100) | Chlorellales(100) | Chlorellales_fa(100) | Planktochlorella(100) |
| ASV 23 | 2485 | Eukaryota(100) | Ciliophora(100) | Intramacronucleata(100) | Conthreep(100) | Oligohymenophorea(100) | Oligohymenophorea_uncl.(100) |
| ASV 24 | 2353 | Eukaryota(100) | Ochrophyta_ph(100) | Chrysophyceae(100) | Ochromonadales(100) | Ochromonadales_fa(100) | Paraphysomonas(100) |
| ASV 25 | 2330 | Eukaryota(100) | Cercozoa(100) | Thecofilosea(100) | Cryomonadida(100) | Rhizaspididae(100) | Rhogostoma(100) |

| | | | | | | | |
|---|---|---|---|---|---|---|---|
| ASV 26 | 2184 | Eukaryota(100) | Diatomea(100) | Bacillariophyceae(100) | Bacillariophyceae(100) | Bacillariophyceae_fa(100) | Nitzschia(100) |
| ASV 27 | 1875 | Eukaryota(100) | Aphelidea(100) | Aphelidea_cl(100) | Aphelidea_or(100) | Aphelidea_fa(100) | uncultured(100) |
| ASV 28 | 1802 | Eukaryota(100) | Ciliophora(100) | Intramacronucleata(100) | Conthreep(100) | Oligohymenophorea(100) | Peritrichia_ge(100) |
| ASV 29 | 1800 | Eukaryota(100) | Ciliophora(100) | Intramacronucleata(100) | Litostomatea(100) | Haptoria(100) | Didinium(100) |
| ASV 30 | 1684 | Eukaryota(100) | Chlorophyta_ph(100) | Chlorophyceae(100) | Sphaeropleales(100) | Sphaeropleales_fa(100) | Sphaeropleales_unclass.(100) |
| ASV 31 | 1606 | Eukaryota(100) | Chlorophyta_ph(100) | Chlorophyceae(100) | Sphaeropleales(100) | Sphaeropleales_fa(100) | Sphaeropleales_unclass.(100) |
| ASV 32 | 1571 | Eukaryota(100) | Ciliophora(100) | Intramacronucleata(100) | Litostomatea(100) | Haptoria(100) | Hemiophrys(100) |
| ASV 33 | 1444 | Eukaryota(100) | Ochrophyta_ph(100) | Chrysophyceae(100) | Ochromonadales(100) | Ochromonadales_fa(100) | Ochromonas(100) |
| ASV 34 | 1355 | Eukaryota(100) | Ciliophora(100) | Intramacronucleata(100) | Litostomatea(100) | Haptoria(100) | Haptoria_unclass.(100) |
| ASV 35 | 1191 | Eukaryota(100) | Eukaryota_ unclass.(100) | Eukaryota_unclass.(100) | Eukaryota_unclass.(100) | Eukaryota_unclass.(100) | Eukaryota_unclass.(100) |
| ASV 36 | 1190 | Eukaryota(100) | Aphelidea(100) | Aphelidea_cl(100) | Aphelidea_or(100) | Aphelidea_fa(100) | Aphelidea_fa_unclass.(100) |
| ASV 37 | 1167 | Eukaryota(100) | Chlorophyta_ph(100) | Trebouxiophyceae(100) | Chlorellales(100) | Chlorellales_fa(100) | Chlorellales_fa_unclass.(100) |
| ASV 38 | 1162 | Eukaryota(100) | Ciliophora(100) | Intramacronucleata(100) | Litostomatea(100) | Haptoria(100) | Loxophyllum(100) |
| ASV 39 | 1107 | Eukaryota(100) | Ciliophora(100) | Intramacronucleata(100) | Conthreep(100) | Oligohymenophorea(100) | Peritrichia_ge(100) |
| ASV 40 | 1046 | Eukaryota(100) | Ciliophora(100) | Intramacronucleata(100) | Conthreep(100) | Oligohymenophorea(100) | Peritrichia_ge(100) |
| ASV 41 | 990 | Eukaryota(100) | Ochrophyta_ph(100) | Chrysophyceae(100) | Ochromonadales(100) | Ochromonadales_fa(100) | Ochromonas(100) |
| ASV 42 | 913 | Eukaryota(100) | Eukaryota_unclass.(100) | Eukaryota_unclass.(100) | Eukaryota_unclass.(100) | Eukaryota_unclassified(100) | Eukaryota_unclass.(100) |
| ASV 43 | 901 | Eukaryota(100) | Chlorophyta_ph(100) | Chlorophyceae(100) | Sphaeropleales(100) | Sphaeropleales_fa(100) | Sphaeropleales_unclass.(100) |
| ASV 44 | 836 | Eukaryota(100) | Ochrophyta_ph(100) | Chrysophyceae(100) | Chrysophyceae(100) | Chrysophyceae_unclass.(100) | Chrysophyceae_unclass.(100) |
| ASV 45 | 827 | Eukaryota(100) | Chlorophyta_ph(100) | Chlorophyceae(100) | Sphaeropleales(100) | Sphaeropleales_fa(100) | Monoraphidium(100) |
| ASV 46 | 813 | Eukaryota(100) | Ciliophora(100) | Intramacronucleata(100) | Conthreep(100) | Oligohymenophorea(100) | Peritrichia_ge(100) |
| ASV 47 | 777 | Eukaryota(100) | Ochrophyta_ph(100) | Chrysophyceae(100) | Ochromonadales(100) | Ochromonadales_fa(100) | Paraphysomonas(100) |
| ASV 48 | 758 | Eukaryota(100) | Ciliophora(100) | Intramacronucleata(100) | Conthreep(100) | Oligohymenophorea(100) | Peritrichia_ge(100) |
| ASV 49 | 738 | Eukaryota(100) | Chlorophyta_ph(100) | Trebouxiophyceae(100) | Chlorellales(100) | Chlorellales_fa(100) | Chlorella(100) |
| ASV 50 | 736 | Eukaryota(100) | Chlorophyta_ph(100) | Chlorophyceae(100) | Sphaeropleales(100) | Sphaeropleales_fa(100) | Desmodesmus(100) |

**Table S4:** Taxonomic information for the top 50 ASVs from 16S rRNA gene sequencing.

| ASV | Size | Taxonomy | | | | | |
|---|---|---|---|---|---|---|---|
| ASV 1 | 551369 | Bacteria(100) | Cyanobacteria(100) | Cyanobacteriia(100) | Cyanobacteriia_unclass(100) | Cyanobacteriia_unclass(100) | Cyanobacteriia_unclass(100) |
| ASV 2 | 176766 | Bacteria(100) | Cyanobacteria(100) | Cyanobacteriia(100) | Cyanobacteriia_unclass(100) | Cyanobacteriia_unclass(100) | Cyanobacteriia_unclass(100) |
| ASV 3 | 119929 | Bacteria(100) | Proteobacteria(100) | Gammaproteobacteria(100) | Burkholderiales(100) | Gallionellaceae(100) | Candidatus_Nitrotoga(100) |
| ASV 4 | 103257 | Bacteria(100) | Cyanobacteria(100) | Cyanobacteriia(100) | Cyanobacteriia_unclass(100) | Cyanobacteriia_unclass(100) | Cyanobacteriia_unclass(100) |
| ASV 5 | 83017 | Bacteria(100) | Cyanobacteria(100) | Cyanobacteriia(100) | Cyanobacteriia_unclass(100) | Cyanobacteriia_unclass(100) | Cyanobacteriia_unclass(100) |
| ASV 6 | 80786 | Bacteria(100) | Bdellovibrionota(100) | Oligoflexia(100) | Oligoflexales(100) | Oligoflexaceae(100) | Oligoflexus(100) |
| ASV 7 | 66846 | Bacteria(100) | Actinobacteriota(100) | Actinobacteria(100) | Corynebacteriales(100) | Mycobacteriaceae(100) | Mycobacterium(100) |
| ASV 8 | 57881 | Bacteria(100) | Proteobacteria(100) | Gammaproteobacteria(100) | Burkholderiales(100) | Nitrosomonadaceae(100) | Nitrosomonas(100) |
| ASV 9 | 40472 | Bacteria(100) | Proteobacteria(100) | Alphaproteobacteria(100) | Sphingomonadales(100) | Sphingomonadaceae(100) | Sphingomonadaceae_unclass(100) |
| ASV 10 | 39629 | Bacteria(100) | Cyanobacteria(100) | Cyanobacteriia(100) | Limnotrichales(100) | Limnotrichaceae(100) | Limnothrix(100) |
| ASV 11 | 35007 | Bacteria(100) | Planctomycetota(100) | Planctomycetes(100) | Planctomycetales(100) | uncultured(100) | uncultured_ge(100) |
| ASV 12 | 30217 | Bacteria(100) | Proteobacteria(100) | Gammaproteobacteria(100) | Burkholderiales(100) | Comamonadaceae(100) | Simplicispira(100) |
| ASV 13 | 28284 | Bacteria(100) | Cyanobacteria(100) | Cyanobacteriia(100) | Cyanobacteriia_unclass(100) | Cyanobacteriia_unclass(100) | Cyanobacteriia_unclass(100) |
| ASV 14 | 26497 | Bacteria(100) | Actinobacteriota(100) | Actinobacteria(100) | Corynebacteriales(100) | Dietziaceae(100) | Dietzia(100) |
| ASV 15 | 25990 | Bacteria(100) | Proteobacteria(100) | Alphaproteobacteria(100) | Rhizobiales(100) | Rhizobiales_Incertae_Sedis(100) | uncultured(100) |
| ASV 16 | 23168 | Bacteria(100) | Proteobacteria(100) | Gammaproteobacteria(100) | Thiotrichales(100) | Thiotrichaceae(100) | Thiothrix(100) |
| ASV 17 | 22761 | Bacteria(100) | Proteobacteria(100) | Gammaproteobacteria(100) | Burkholderiales(100) | Nitrosomonadaceae(100) | Nitrosomonas(100) |
| ASV 18 | 22533 | Bacteria(100) | Chloroflexi(100) | Anaerolineae(100) | RBG-13-54-9(100) | RBG-13-54-9_fa(100) | RBG-13-54-9_ge(100) |
| ASV 19 | 20131 | Bacteria(100) | Planctomycetota(100) | Planctomycetes(100) | Isosphaerales(100) | Isosphaeraceae(100) | uncultured(100) |
| ASV 20 | 18796 | Bacteria(100) | Proteobacteria(100) | Gammaproteobacteria(100) | Burkholderiales(100) | Nitrosomonadaceae(100) | Nitrosomonas(100) |
| ASV 21 | 18032 | Bacteria(100) | Verrucomicrobiota(100) | Verrucomicrobiae(100) | Chthoniobacterales(100) | Chthoniobacteraceae(100) | Chthoniobacter(100) |
| ASV 22 | 17851 | Bacteria(100) | Proteobacteria(100) | Gammaproteobacteria(100) | Burkholderiales(100) | Gallionellaceae(100) | Candidatus_Nitrotoga(100) |
| ASV 23 | 17161 | Bacteria(100) | Planctomycetota(100) | Planctomycetes(100) | Planctomycetales(100) | Rubinisphaeraceae(100) | SH-PL14(100) |
| ASV 24 | 17159 | Bacteria(100) | Proteobacteria(100) | Gammaproteobacteria(100) | Xanthomonadales(100) | Xanthomonadaceae(100) | Xanthomonadaceae_unclass(100) |
| ASV 25 | 17118 | Bacteria(100) | Proteobacteria(100) | Gammaproteobacteria(100) | Burkholderiales(100) | Nitrosomonadaceae(100) | Nitrosomonas(100) |

| | | | | | | | |
|---|---|---|---|---|---|---|---|
| ASV 26 | 16174 | Bacteria(100) | Planctomycetota(100) | Planctomycetes(100) | Pirellulales(100) | Pirellulaceae(100) | Pirellula(100) |
| ASV 27 | 15924 | Bacteria(100) | Acidobacteriota(100) | Blastocatellia(100) | 11-24(100) | 11-24_fa(100) | 11-24_ge(100) |
| ASV 28 | 15210 | Bacteria(100) | Firmicutes(100) | Clostridia(100) | Peptostreptococcales-Tissierellales(100) | Peptostreptococcaceae(100) | Romboutsia(100) |
| ASV 29 | 14980 | Bacteria(100) | Proteobacteria(100) | Gammaproteobacteria(100) | Xanthomonadales(100) | Rhodanobacteraceae(100) | Rhodanobacter(100) |
| ASV 30 | 14288 | Bacteria(100) | Bacteroidota(100) | Bacteroidia(100) | Cytophagales(100) | Microscillaceae(100) | OLB12(100) |
| ASV 31 | 13531 | Bacteria(100) | Actinobacteriota(100) | Actinobacteria(100) | Corynebacteriales(100) | Mycobacteriaceae(100) | Mycobacterium(100) |
| ASV 32 | 12901 | Bacteria(100) | Proteobacteria(100) | Gammaproteobacteria(100) | Burkholderiales(100) | Comamonadaceae(100) | Comamonadaceae_unclass(100) |
| ASV 33 | 12569 | Bacteria(100) | Bacteroidota(100) | Bacteroidia(100) | Flavobacteriales(100) | Flavobacteriaceae(100) | Flavobacterium(100) |
| ASV 34 | 12081 | Bacteria(100) | Planctomycetota(100) | Planctomycetes(100) | Planctomycetales(100) | uncultured(100) | uncultured_ge(100) |
| ASV 35 | 11955 | Bacteria(100) | Bacteroidota(100) | Bacteroidia(100) | Bacteroidia_unclass(100) | Bacteroidia_unclassified(100) | Bacteroidia_unclass(100) |
| ASV 36 | 11376 | Bacteria(100) | Proteobacteria(100) | Gammaproteobacteria(100) | Burkholderiales(100) | Comamonadaceae(100) | Rhodoferax(100) |
| ASV 37 | 11107 | Bacteria(100) | Firmicutes(100) | Bacilli(100) | Lactobacillales(100) | Carnobacteriaceae(100) | Trichococcus(100) |
| ASV 38 | 10894 | Bacteria(100) | Chloroflexi(100) | Anaerolineae(100) | Caldilineales(100) | Caldilineaceae(100) | uncultured(100) |
| ASV 39 | 10622 | Bacteria(100) | Bacteroidota(100) | Bacteroidia(100) | Chitinophagales(100) | Saprospiraceae(100) | uncultured(100) |
| ASV 40 | 10462 | Bacteria(100) | Proteobacteria(100) | Alphaproteobacteria(100) | Rhodobacterales(100) | Rhodobacteraceae(100) | Rhodobacter(100) |
| ASV 41 | 10453 | Bacteria(100) | Nitrospirota(100) | Nitrospiria(100) | Nitrospirales(100) | Nitrospiraceae(100) | Nitrospira(100) |
| ASV 42 | 10394 | Bacteria(100) | Actinobacteriota(100) | Actinobacteria(100) | Corynebacteriales(100) | Mycobacteriaceae(100) | Mycobacterium(100) |
| ASV 43 | 10295 | Bacteria(100) | Planctomycetota(100) | Planctomycetes(100) | Planctomycetales(100) | Rubinisphaeraceae(100) | uncultured(100) |
| ASV 44 | 10288 | Bacteria(100) | Proteobacteria(100) | Alphaproteobacteria(100) | Rhizobiales(100) | Devosiaceae(100) | Devosia(100) |
| ASV 45 | 10163 | Bacteria(100) | Actinobacteriota(100) | Actinobacteria(100) | Corynebacteriales(100) | Mycobacteriaceae(100) | Mycobacterium(100) |
| ASV 46 | 9768 | Bacteria(100) | Planctomycetota(100) | Planctomycetes(100) | Pirellulales(100) | Pirellulaceae(100) | Pirellula(100) |
| ASV 47 | 9346 | Bacteria(100) | Proteobacteria(100) | Alphaproteobacteria(100) | Sphingomonadales(100) | Sphingomonadaceae(100) | Sphingopyxis(100) |
| ASV 48 | 8885 | Bacteria(100) | Proteobacteria(100) | Gammaproteobacteria(100) | Xanthomonadales(100) | Rhodanobacteraceae(100) | Dokdonella(100) |
| ASV 49 | 8837 | Bacteria(100) | Proteobacteria(100) | Gammaproteobacteria(100) | Burkholderiales(100) | Rhodocyclaceae(100) | Rhodocyclaceae_unclass(100) |
| ASV 50 | 8613 | Bacteria(100) | Proteobacteria(100) | Alphaproteobacteria(100) | Sphingomonadales(100) | Sphingomonadaceae(100) | Rhizorhapis(100) |

Lee, P.A., Martinez, K.J.L., Letcher, P.M., Corcoran, A.A., Ryan, R.A., 2018. A novel predatory bacterium infecting the eukaryotic alga Nannochloropsis. Algal Res 32, 314–320. https://doi.org/10.1016/J.ALGAL.2018.04.003
Waite, D.W., Chuvochina, M., Pelikan, C., Parks, D.H., Yilmaz, P., Wagner, M., Loy, A., Naganuma, T., Nakai, R., Whitman, W.B., Hahn, M.W., Kuever, J., Hugenholtz, P., 2020. Proposal to reclassify the proteobacterial classes Deltaproteobacteria and Oligoflexia, and the phylum Thermodesulfobacteria into four phyla reflecting major functional capabilities. Int. J. Syst. Evol. Microbiol 70, 5972–6016. https://doi.org/10.1099/ijsem.0.004213
